# Identifying loci under positive selection in complex population histories

**DOI:** 10.1101/453092

**Authors:** Alba Refoyo-Martínez, Rute R. da Fonseca, Katrín Halldórsdóttir, Einar Árnason, Thomas Mailund, Fernando Racimo

## Abstract

Detailed modeling of a species’ history is of prime importance for understanding how natural selection operates over time. Most methods designed to detect positive selection along sequenced genomes, however, use simplified representations of past histories as null models of genetic drift. Here, we present the first method that can detect signatures of strong local adaptation across the genome using arbitrarily complex admixture graphs, which are typically used to describe the history of past divergence and admixture events among any number of populations. The method—called Graph-aware Retrieval of Selective Sweeps (*GRoSS*)—has good power to detect loci in the genome with strong evidence for past selective sweeps and can also identify which branch of the graph was most affected by the sweep. As evidence of its utility, we apply the method to bovine, codfish and human population genomic data containing multiple population panels related in complex ways. We find new candidate genes for important adaptive functions, including immunity and metabolism in under-studied human populations, as well as muscle mass, milk production and tameness in specific bovine breeds. We are also able to pinpoint the emergence of large regions of differentiation due to inversions in the history of Atlantic codfish.

## Introduction

One of the main goals of population genomics is to understand how adaptation affects patterns of variation across the genome and to find ways to analyze these patterns. In order to identify loci that have been affected by positive selection in the past, geneticists have developed methods that can scan a set of genomes for signals that are characteristic of this process. These signals may be based on patterns of haplotype homozygosity [1, 2], the site frequency spectrum [3, 4] or allelic differentiation between populations [5, 6].

Population differentiation-based methods have proven particularly successful in recent years, as they make few assumptions about the underlying demographic process that may have generated a selection signal, and are generally more robust and scalable to large population-wide datasets. The oldest of these are based on computing pairwise *F_ST_* [7, 8] or similar measures of population differentiation between two population panels across SNPs or windows of the genome [9, 10]. More recent methods have allowed researchers to efficiently detect which populations are affected by a sweep, by computing branch-specific differentiation on 3-population trees [6, 11, 12], 4-population trees [13], or arbitrarily large population trees [14–16], or by looking for strong locus-specific differentiation or environmental correlations, using the genome-wide population-covariance matrix as a null model of genetic drift [17–21].

Although some of these methods for detecting selection implicitly handle past episodes of admixture, none of them uses “admixture graphs” that explicitly model both divergence and admixture in an easily interpretable framework [22, 23]. This makes it difficult to understand the signatures of selection when working with sets of multiple populations that may be related to each other in complex ways. Here, we introduce a method to efficiently detect selective sweeps across the genome when using many populations that are related via an arbitrarily complex history of population splits and mergers. We modified the *Q_B_* statistic [24] which was originally meant to detect polygenic adaptation using admixture graphs. Unlike *Q_B_*, our new statistic—which we call *S_B_*—does not need gene-trait association data and works with allele frequency data alone. It can be used to both scan the genome for regions under strong single-locus positive selection, and to pinpoint where in the population graph the selective event most likely took place. We demonstrate the usefulness of this statistic by performing selection scans on human, bovine and codfish data, recovering existing and new candidate loci, while obtaining a clear picture of which populations were most affected by positive selection in the past.

## Methods

### Theory

We modified the previously-developed *Q_B_* statistic [24] to detect strong branch-specific deviations in single-locus allele frequencies, but without having to use effect size estimates from an association study. We assume that the topology of the admixture graph relating a set of populations is known and that we have allele frequency data for all the populations we are studying. For a single SNP, let **p** be the vector of allele frequencies across populations. We then make a multivariate normal approximation to obtain a distribution with which we can model these frequencies [14, 17, 18, 25]:

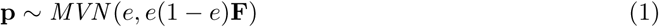

where F is the neutral covariance matrix and e is the ancestral allele frequency of all populations. We use the genome-wide covariance matrix as an estimate of the neutral covariance matrix. In practice, the ancestral allele frequency is unknown, so the mean allele frequency among populations can be used as an approximate stand-in. We then obtain a mean-centered version of the vector **p**, which we call **y**:

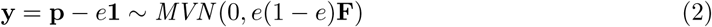

where **1** is a column vector of as many ones as there are populations. For an arbitrarily-defined, mean-centered vector **b** with the same number of elements as there are populations:

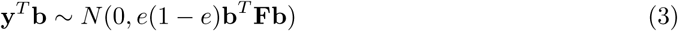

Our test statistic—which we call *S_B_*—is then defined as:

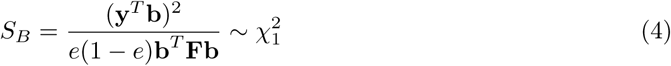

The key is to choose a vector **b** that represents a specific branch of our graph. Essentially, for a branch *j*, the elements of its corresponding branch vector **b_j_** are the ancestry contributions of that branch to each of the populations in the leaves of the graph. For a more detailed description of how to construct this vector, see Racimo, Berg and Pickrell [24].

If we choose **b** to be the vector corresponding to branch *j* when computing the statistic in equation 4, then significant values of the statistic *S_B_*(*j*) will capture deviations from neutrality in the graph that are attributable to a disruption that occurred along branch *j*.

If we only have a few genomes per population, the true population allele frequencies will be poorly estimated by our sample allele frequencies, potentially decreasing power. However, we can increase power at the cost of spatial genomic resolution and rigorous statistical interpretation, by combining information from several SNPs into windows, as was done, for example, in Skoglund et al. [26]. We can compute the average χ^2^ statistic over all SNPs in each window and provide a new P-value for that averaged statistic. As the chi-squared distributional assumption only holds for low amounts of drift, it may be useful to standardize the scores using the mean and variance of the genome-wide distribution, especially when working with populations that diverged from each other a long time ago.

### Implementation

We implemented the *S_B_* statistic in a program called *GRoSS* (Graph-aware Retrieval of Selective Sweeps), which is freely available on GitHub: https://github.com/FerRacimo/GRoSS. The program runs on R and makes use of the admixturegraph R library [27]. We also wrote a module that allows one to input a file specifying the admixture graph topology directly.

Figure 1 shows a schematic workflow for *GRoSS*. The user begins by estimating an admixture graph using genome-wide data, via a program like *TreeMix* [23], *MixMapper* [28] or *qpGraph* [22]. Then, the user writes the topology of the graph to a text file. The format of this file can be either the dot-format or the input file format for *qpGraph*, so it can be skipped if the initial step was run using *qpGraph*. Then, the user inputs the graph topology, and a file with major/minor allele counts for each SNP into *GRoSS*. The allele counts can also be polarized as ancestral/derived or reference/alternative. *GRoSS* will compute the genome-wide covariance matrix and the **b** vectors for each branch, and then calculate the *S_B_* scores and corresponding **P**-values, which can then be plotted.

**Figure 1:**
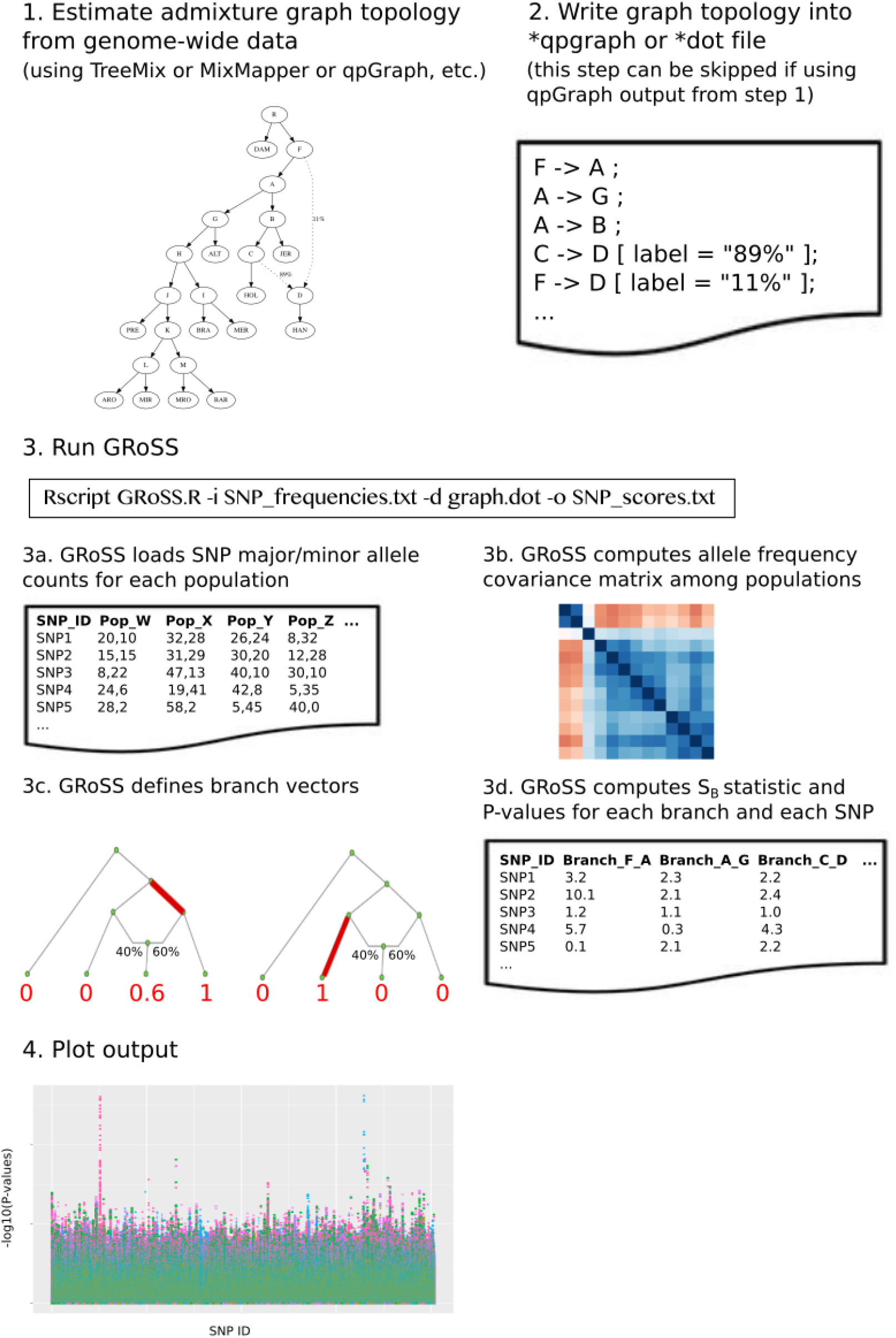
Schematic of *GRoSS* workflow

### Simulations

We used SLiM 2 [29] to simulate genomic data and test how our method performs at detecting positive selection, with sample sizes of 100, 50, 25 and 4 diploid genomes per population (Figures 2, S1, S2 and S3, respectively). We simulated a genomic region of length 10Mb, a constant effective population size (*N_e_*) of 10,000, a mutation rate of 10^−8^ per base-pair per generation and a uniform recombination rate of 10^−8^ per base-pair per generation. We placed the beneficial mutation in the middle of the region, at position 5Mb. We used a burn-in period of 100,000 generations to generate steady-state neutral variation. For each demographic scenario that we tested, we simulated under neutrality and two selective regimes, with selection coefficients (s) of 0.1 and 0.01. We considered two types of selection scenarios for each demographic scenario: one in which we condition on the beneficial mutation reaching > 1% frequency at the final generation of the branch in which we simulated the positive selection event, and one in which we condition the mutation reaching > 5% frequency. We discarded simulations that did not fulfill these conditions. We set the time intervals between population splits at 2,000 generations for all branches of the population graph in the 3-population and 6-population graphs, and at 500 generations in the 16-population graph. To speed up the simulations, we scaled the values of the population size and of time by a factor of 1/10 and, consequently, the mutation rate, recombination rate and selection coefficients by a factor of 10 [29].

**Figure 2:**
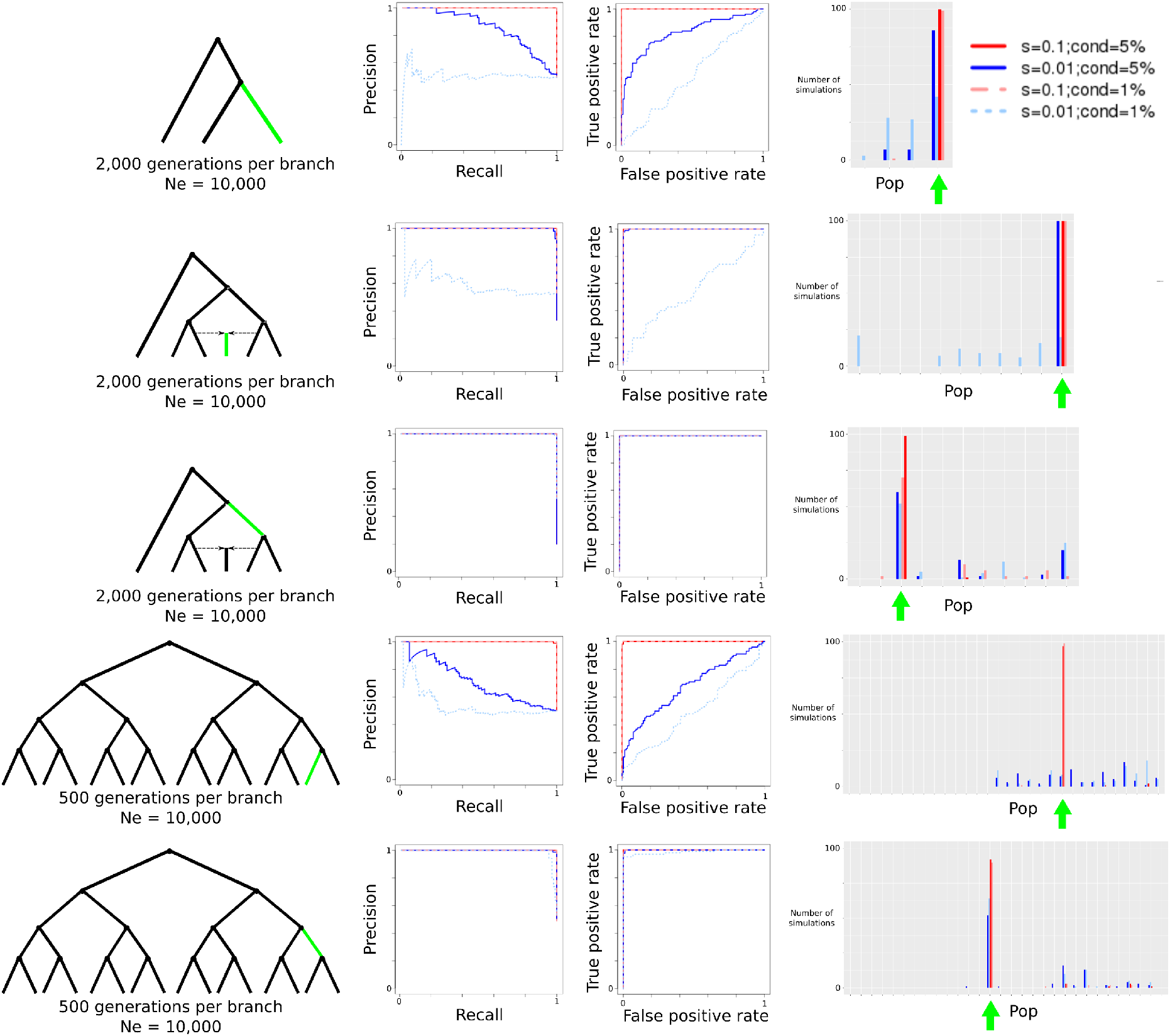
Evaluation of *GRoSS* performance using simulations in *SLiM 2*, with 100 diploid individuals per population panel. We simulated different selective sweeps under strong (s=0.1) and intermediate (s=0.01) selection coefficients for a 3-population tree, a 6-population graph with a 50%/50% admixture event and a 16-population tree. We then produced precision-recall and ROC curves comparing simulations under selection to simulations under neutrality. We also obtained the maximum branch score within 100kb of the selected site, and computed the number of simulations (out of 100) in which the branch of this score corresponded to the true selected branch. “*cond* = 5%”: Simulations conditional on the beneficial mutation reaching 5% frequency or more. “*cond* = 1%”: Simulations conditional on the beneficial mutation reaching 1% frequency or more. “Pop”: population branch.

### Human data

We used data from Phase 3 of the 1000 Genomes Project [30] and a SNP chip dataset of present-day humans from 203 populations genotyped with the Human Origins array [22, 31]. The SNP chip dataset was imputed using *SHAPEIT* [32] on the Michigan Imputation Server [33] with the 1000 Genomes data as the reference panel [24]. We used inferred admixture graphs that were fitted to this panel using *MixMapper* (v1.02) [28] in a previous publication [24]. For the 1000 Genomes dataset, the inferred graph was a tree where the leaves are composed of panels from 7 populations: Southern Han (CDX), Han Chinese from Beijing (CHB), Japanese from Tuscany (JPT), Toscani (TSI), Utah Residents (CEPH) with Northern and Western European Ancestry (CEU), Mende from Sierra Leone (MSL) and Esan from Nigeria (ESN) (Figure 3). For the Human Origins dataset, the inferred graph was a 7-leaf admixture graph that includes Native Americans, East Asians, Oceanians, Mandenka, Yoruba, Sardinians and Europeans with high ancient-steppe (Yamnaya) ancestry (Figure 3). This graph contains an admixture event from a sister branch to Native Americans and a sister branch to Sardinians into Europeans, representing the ancient steppe ancestry known to be present in almost all present-day Europeans (but largely absent in present-day Sardinians).

**Figure 3:**
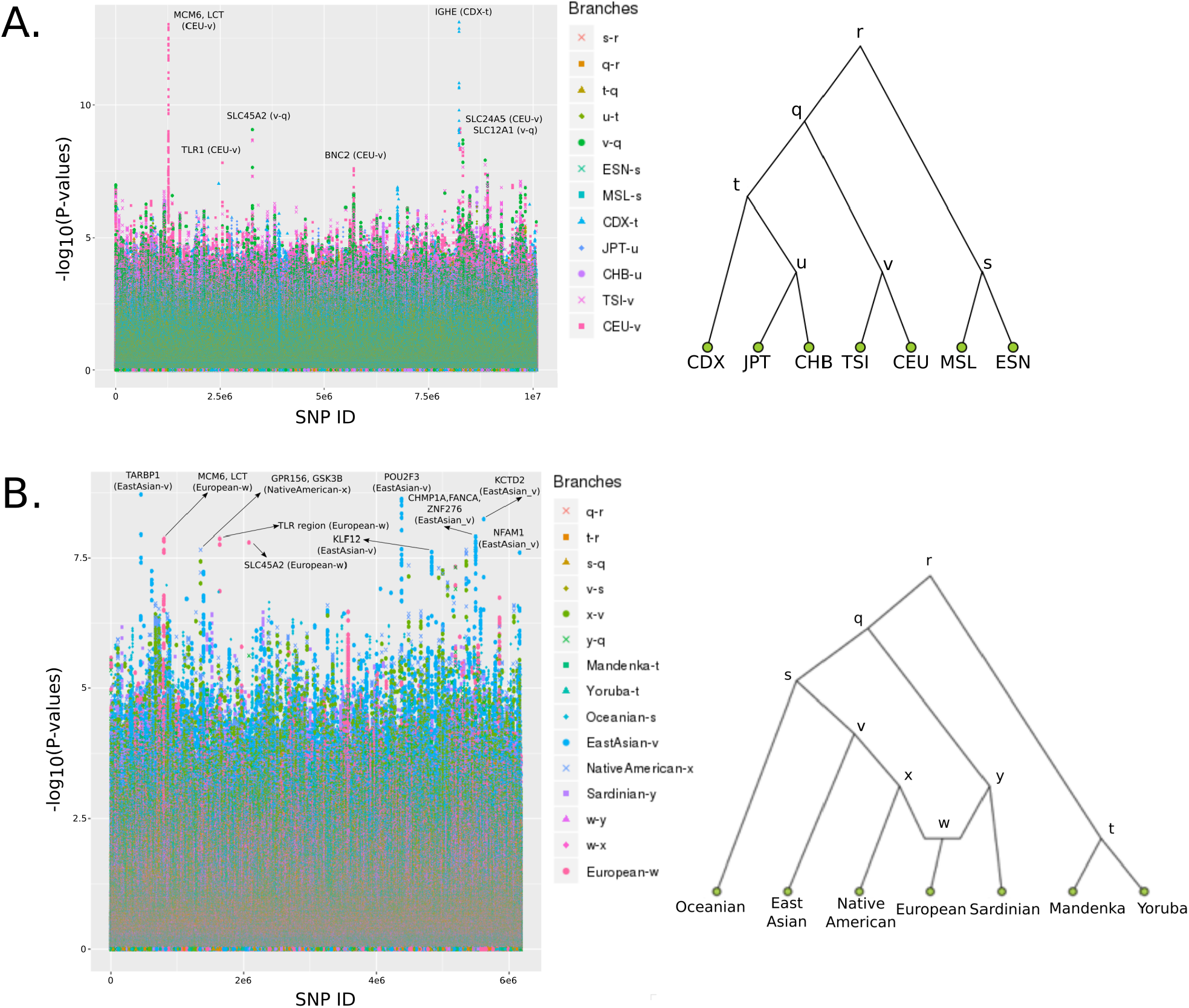
Results from running *GRoSS* on human genomic data. A) Population tree including panels from phase 3 of the 1000 Genomes project. B) Population graph including imputed panels from the Human Origins SNP chip data from Lazaridis et al. (2014).

We removed sites with < 1% minor allele frequency or where at least one population had no coverage. We then ran *GRoSS* on the resulting SNPs in each of the two datasets (Tables S1 and S2). We selected SNPs with –*log*_10_(*P*) larger than 7, and merged SNPs into regions if they were within 100kb of each other. Finally, we retrieved all HGNC protein-coding genes that overlap each region, using *biomaRt* [34].

### Bovine data

We assembled a population genomic dataset (Table S3) containing different breeds of *Bos taurus* using: a) SNP array data from ref. [35], corresponding to the Illumina BovineHD Genotyping BeadChip (http://dx.doi.org/10.5061/dryad.f2d1q); b) whole-genome shotgun data from 10 individuals from the indigenous African breed NĎama [36] (Bioproject ID: PRJNA312138); c) shotgun data from two commercial cattle breeds (Holstein and Jersey; Bioproject IDs: PRJNA210521 and PRJNA318089, respectively); d) shotgun data for 8 Iberian cattle breeds [37].

We used *TreeMix* [23] to infer an admixture graph (Figure S4) using allele counts for 512,358 SNPs in positions that were unambiguously assigned to the autosomes in the cattle reference genome version UMD_3.1.1 [38] using SNPchiMp [39]. For shotgun data, allele counts were obtained from allele frequencies calculated in *ANGSD* [40] for positions covered in at least three individuals. We removed SNPs for which at least one panel had no coverage or in which the minor allele frequency was less than 1%.

We applied the statistic to the *TreeMix*-fitted graph model in Figure 4. We performed the scan in two ways: in one we computed a per-SNP chi-squared statistic, from which we obtained a P-value (Table S4), and in the other we combined the chi-squared statistics in windows of 10 SNPs (Table S5), with a step size of 1 SNP, obtaining a P-value for a particular window using its average *S_B_* score (Figure 4). We used this windowing scheme because of concerns about small sample sizes in some of the populations, and aimed to pool information across SNPs within a region. After both scans, we combined windows that were within 100kb of each other into larger regions, and retrieved HGNC and VGNC genes within a +/- 100kb window around the boundaries of each region using biomaRt [34] with the April 2018 version of Ensembl.

**Figure 4:**
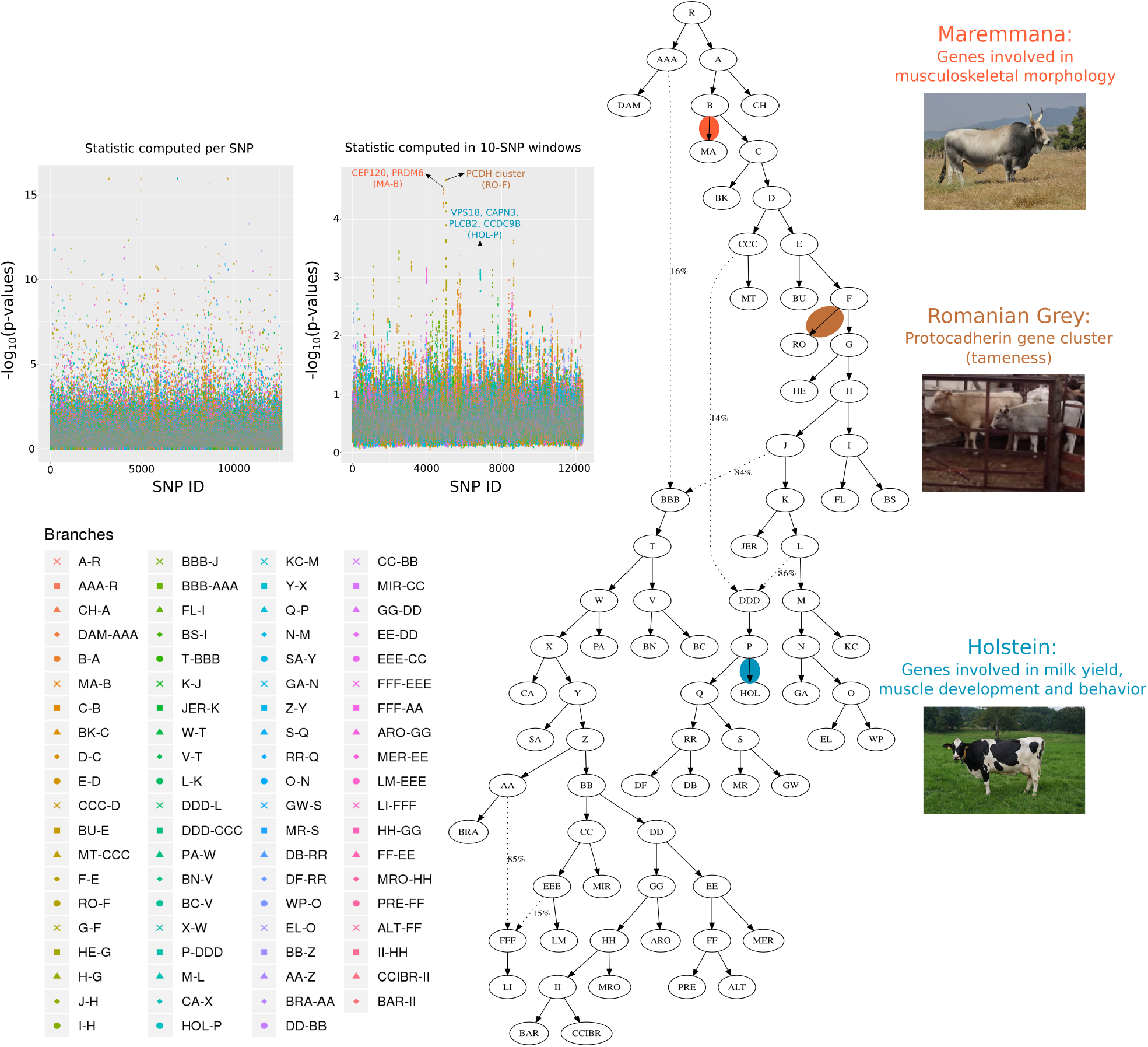
Results from running *GRoSS* on a population graph of bovine breeds. P-values were obtained by either computing chi-squared statistics per SNP, or after averaging the per-SNP statistics in 10-SNP windows with a 1-SNP step size, and obtaining a P-value from the averaged statistic. Holstein and Maremmana cattle photos obtained from Wikimedia Commons (authors: Verum; giovanni bidi). Romanian Grey cattle screen-shot obtained from a CC BY YouTube video (author: Paolo Caddeo).

### Codfish data

Codfish were randomly sampled from a large tissue sample database [41] and the J. Mork collections from populations covering a wide distribution from the western Atlantic to the northern and eastern Atlantic (Figure S5 and Table S6). The populations differ in various life-history and other biological traits [42, 43], and their local environment ranges from shallow coastal water (e.g. western Atlantic and North Sea) to waters of great depth (e.g. parts of Iceland and Barents Sea). They also differ in temperature and salinity (e.g. brackish water in the Baltic).

We isolated genomic DNA from gill tissue and fin clips stored in ethanol using the E.Z.N.A.^®^ Tissue DNA Kit (Omega biotek) following the manufacturer’s protocol. Libraries were prepared and individually indexed for sequencing using the Nextera^®^ DNA Library Preparation Kit (Illumina, FC-121–1031). Pooled libraries were sequenced on the HiSeq 2500 in rapid run mode (paired-end, 2×250 cycles) at the Bauer Core Facility at Harvard University. We aligned fastq files to the gadmor2 assembly [44] using *bwa mem* [45], merged and deduplicated using *Picard* (http://broadinstitute.github.io/picard) and *GATK* [46] in accordance with *GATK* best practices [47]. Details of the molecular and analysis methods are given in [48, 49]. We ran *ANGSD* [40] on the genome sequences from all populations, computed base-alignment quality [50], adjusted mapping quality for excessive mismatches, and filtered for mapping quality (≥ 30) and base quality (≥ 20). We then estimated the allele frequencies in each population at segregating sites using the –`sites` option of *ANGSD*.

We applied the statistic to the graph model in Figure 6, estimated using *TreeMix* [23] allowing for 3 migration events (Figure S6). We removed SNPs where at least 1 panel had no coverage or in which the minor allele frequency was less than 1%, and we only selected sites in which all panels had 2 or more diploid individuals covered. We performed the scan by combining the per-SNP chi-squared statistics in windows of 10 SNPs, with a step size of 5 SNPs, obtaining a P-value for a particular window using its average *S_B_* score (Figures S7, S8, S9, S10). In a preliminary analysis, we identified 4 large regions of high differentiation related to structural variants, which span several mega-bases (see Results and Discussion below). In our final analysis, we excluded sites lying within linkage groups that contain these regions from the *TreeMix*-fitting and covariance matrix estimation, so as to prevent them from biasing our null genome-wide model.

**Figure 5:**
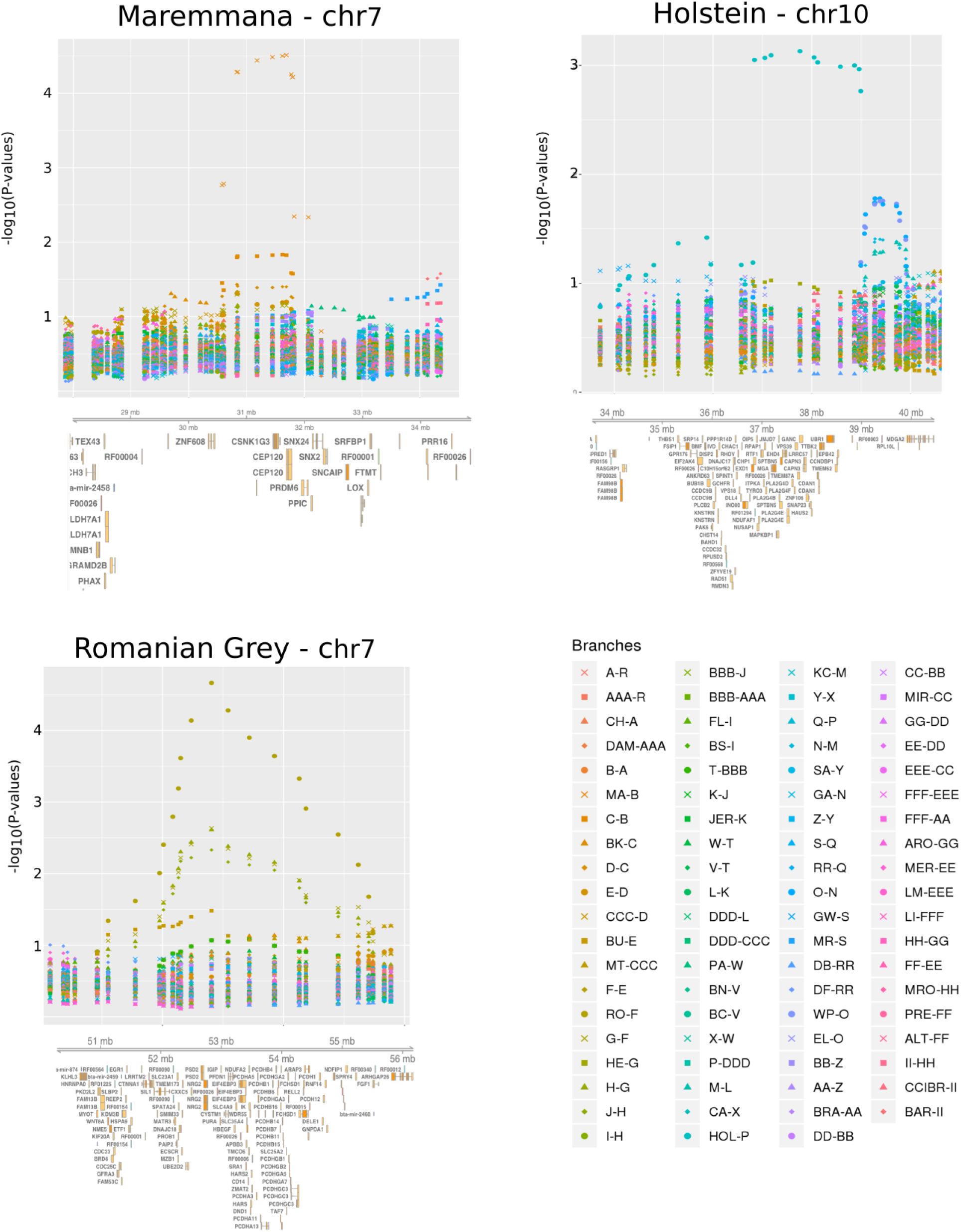
Zoomed-in plots of GRoSS output for three regions found to have strong evidence for positive selection in the 10-SNP bovine scan. Genes were retrieved using Ensembl via the Gviz R Bioconductor library [96].

**Figure 6:**
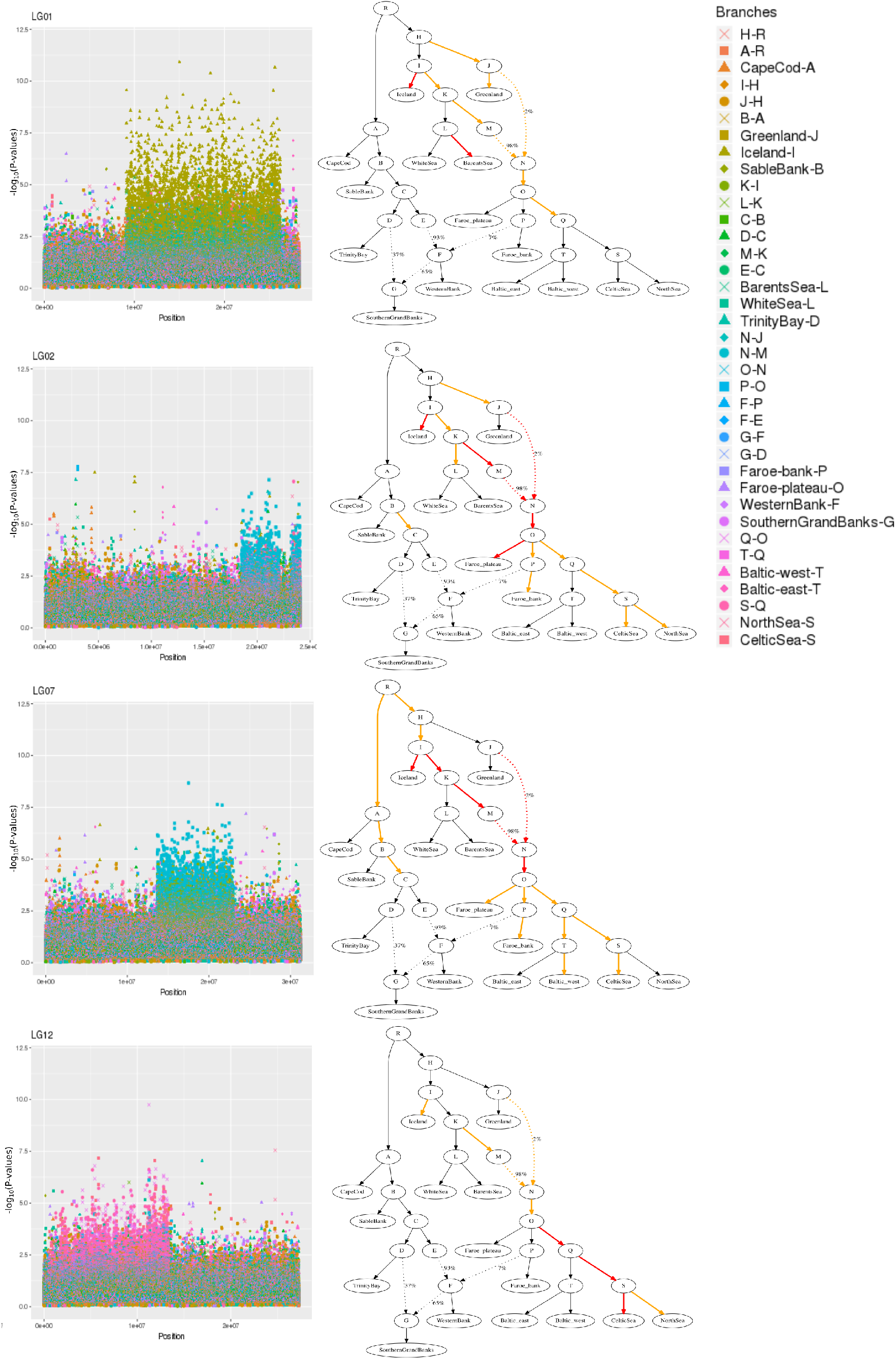
Large regions of high differentiation in the codfish data. Branches colored in orange are branches whose corresponding *S_B_* scores evince the high-differentiation region. Branches colored in red are branches whose corresponding *S_B_* scores evince the high-differentiation region and have at least one SNP with –*log*_10_(*P*) > 5 inside the region.

### Selection of candidate regions

Given the myriad of plausible violations of our null multivariate-Normal model (see Discussion), we do not expect the P-values of the *S_B_* statistic to truly reflect the probability one has of rejecting a neutral model of evolution. We therefore see these P-values as a guideline for selecting regions as candidates for positive selection, rather than a way for rigorously determining the probability that a region has been evolving neutrally. In all applications below, we used arbitrary P-value cutoffs to select the top candidate regions based on visual inspection of the Manhattan plots of the genome-wide distribution of *S_B_* P-values. These empirical cutoffs vary across study species and also depend on the specific scheme we use to calculate the *S_B_* statistic (per-SNP or averaged over a window) and we do not claim these cutoffs to have any statistical motivation beyond being convenient ways to separate regions that lie at the tails of our empirical distribution.

Alternative approaches could involve using a randomization scheme or generating simulations based on a fitted demographic model to obtain a neutral distribution of loci and derive a P-value from that. While any of those approaches could be pursued with the *S_B_* framework, we do not pursue any of those approaches in this paper. We think that the chosen mode of randomization or the fitted demographic parameters will also necessarily rely on assumptions about unknown or unmodeled parameters, and may provide unmerited confidence to the cutoff that we could end up choosing. Instead, we recommend that the reader take our chi-squared-distributed P-values with a grain of salt, and merely use them as a way to prioritize regions for more extensive downstream modeling and validation (for example, using methods like those described in refs. [51–53]).

## Results

### Simulations

We performed simulations on SLiM 2 [29], and used ROC and precision-recall curves to evaluate the performance of our method under different demographic scenarios, and to compare the behavior of our scores under selection and neutrality. For each demographic scenario, we tested four selective sweep modes: comparing simulations under two different selection coefficients (s=0.1 and s=0.01) against a set of neutral simulations (s=0), and, for each of these, conditioning on establishment of the beneficial mutation at more than 5% frequency or at more than 1% frequency. Each branch of each graph had a diploid population size (*N_e_*) of 10,000.

First, we simulated an episode of positive selection occurring on a branch of a three-population tree with no admixture. Each branch of the tree lasted for 2,000 generations. We sampled 100 individuals from each population. Unsurprisingly, the performance of the method under both selection coefficients is higher when we condition on a higher frequency of establishment of the beneficial allele, and is also better under strong selection (Figure 2). We also kept track of which branch in each simulation had the highest score in a region of 100kb centered on the beneficial mutation. As shown in Figure 2, the highest values typically correspond to the population in which selection was simulated.

We then simulated more complex demographic histories including a 6-population graph with admixture. Each branch of the graph lasted for 2,000 generations. We explored two different selection scenarios. In one scenario, the selective sweep was introduced in one of the internal branches whereas in another scenario, it was introduced in one of the external branches. Interestingly, the performance under this graph appears to improve relative to the three-population scenario (Figure 2). The reason is that the *S_B_* statistic depends on having an accurate estimate of the ancestral allele frequency (*e*). This estimate is calculated by taking the average of all allele frequencies in the leaf populations, so the more leaf populations in a well-balanced graph we have, the more accurate this estimate will be.

In Figure 2, we also show the maximum scores for each branch in a 100kb region around the beneficial allele under both scenarios. We find that the branch where the selective sweep was simulated also tends to have the highest *S_B_* scores. We also explored a larger population tree with sixteen leaf populations. In this case, each branch of the graph lasted for 500 generations. ROC and recall curves show a similar performance to the ones from the 6-population admixture graph (Figure 2).

Finally, we explored the performance of the method when the number of diploid individuals per population was smaller than 100. Figure S1 shows the performance of the method with 50 diploid individuals per population, figure S2 shows the performance with 20 individuals and figure S3 shows the performance with 4 individuals. Even when the number of individuals is this small, we can still recover most of the simulated sweeps, especially when selection is strong.

### Positive selection in human populations across the world

Applying our method to both the 1000 Genomes and the Human Origins population graphs, we observe many candidate loci that have been identified in previous world-wide positive selection scans (some of them due to archaic adaptive introgression). Previously reported selection candidates that we recover include the LCT/MCM6, BNC2, OCA2/HERC2, TLR and SLC24A5 regions in northern Europeans, the CHMP1A/ZNF276/FANCA, ABCC11 and POU2F3 regions in East Asians, and the SLC45A2 and SLC12A1 genes in an ancestral European population [1, 11, 12, 54–59] (Tables S1, S2).

We find that the IGH immune gene cluster (also containing gene KIAA0125) is the strongest candidate for selection in the 1000 Genomes scan, and the signal is concentrated on the Chinese Dai branch. This cluster has been recently reported as being under selection in a large Chinese cohort of over 140,000 genomes [60]. Our results suggest that the selective pressures may have existed somewhere in southern China, as we do not see such a strong signal in other parts of the East Asian portion of the graph.

A region containing TARBP1 was the strongest candidate for selection in the Human Origins scan (East Asian terminal branch). The gene codes for an HIV-binding protein and has been previously reported to be under balancing selection [61]. The top SNP (rs2175591) lies in an H3K27Ac regulatory mark upstream of the gene. The derived allele at this SNP is at more than 50% frequency in all 1000 Genomes East Asian panels but at less than 2% frequency in all the other worldwide panels, except for South Asians where it reaches frequencies of around 10%. Interestingly, the TARBP1 gene has been identified as a target for positive selection in milk-producing cattle [62] and in sheep breeds [63, 64]. It has also been associated with resistance to gastrointestinal nematodes in sheep [65]. Our results suggest it may have also played an important role during human evolution in eastern Eurasia, possibly as a response to local pathogens.

Another candidate for selection is the NFAM1 gene in East Asians, which codes for a membrane receptor that is involved in development and signaling of B-cells [66]. This gene was also found to be under positive selection in the Sheko cattle of Ethiopia, along with other genes related to immunity [67].

In the Native American terminal branch of the Human Origins scan, we find a candidate region containing two genes: GPR156 and GSK3B. GPR156 codes for a G protein-coupled receptor, while GSK3B codes for a kinase that plays important roles in neuronal development, energy metabolism, and body pattern formation [68]. We also find a candidate region in the same branch in the protamine gene cluster (PRM1, PRM2, PRM3, TNP2), involved in spermatogenesis [69, 70], and another region overlapping MDGA2, which is specifically expressed in the nervous system [71].

### Cattle breeding: morphology, tameness and milk yield

We performed two scans on the bovine data, one in which computed the S_B_ statistic per SNP (Table S4), and one in which we computed it in 10-SNP windows (Table S5). The window-based scan retrieved 12 top candidate regions, 10 of which overlap with regions previously detected to be under selection in cattle (reviewed in [72]). Additionally, 28 of the 43 top candidate SNPs from the single-SNP scan are also in regions that have been previously reported as selection candidates.

The two top scoring regions from the 10-SNP scan are both in chromosome 7, and include regions that have been detected to be under selection in both taurine and zebu cattle (the latter not represented in our sample), and are potentially associated with cattle traits of general interest in domesticated species. The region located between 30 and 33 Mb of chromosome 7 appears as a top candidate in both the 1-SNP and 10-SNP scan, on the terminal branch leading to the Maremmana (MA) breed (Figure 5). It includes genes related to variation in body shape, such as CEP120 whose mutations have been linked to a type of skeletal dysplasia that results in thoracic cage and extra-skeletal abnormalities [73], and PRDM6, a histone modifier that can induce various smooth muscle phenotypes [74].

The region located between 50 and 55 Mb contains members of the three protocadherin (Pcdha, Pcdhb and Pcdhg) gene clusters (Figure 5). It is identified by *GRoSS* to be under selection in Romanian Grey cattle (RO), which is well-known for its docile disposition. Protocadherins are cell-adhesion molecules that are differentially expressed in individual neurons [75]. They have been implicated in mental retardation and epilepsy in humans [76] and in fear-conditioning and memory in mice [77], and have also been shown to be under selection in cats [78]. Genes of the protocadherin family have also been detected to have expression and allele frequency differences consistent with adaptation in an analysis of tame and aggressive foxes [79].

The largest window (4.4 Mb) detected by GRoSS corresponds to the branch leading to the Holstein (HOL) breed (Figure 5). This window overlaps regions found to be under selection in Holstein using various tests (reviewed in [72]). Some of the outlier genes that were also identified in an earlier XP-EHH scan [80] include VPS18, implicated in neurodegeneration [81] and CAPN3, associated with muscle dystrophy. The window also contains genes that are differentially expressed between high and low milk yield cows (PLCB2 and CCDC9B). [82].

### Large regions of extreme differentiation in Atlantic codfish

When running *GRoSS* on the Atlantic codfish data, we found 4 large genomic regions of high differentiation spanning several mega-bases, on 4 different linkage groups: LG01, LG02, LG07, and LG12 (Figure 6, Table S7). These regions were previously detected by pairwise F_ST_ analyses [48, 83, 84]. They are associated with inversions that suppress recombination in heterozygous individuals, and have thereby favored dramatic increases in differentiation between haplotypes. The signals in the LG01, LG02 and LG07 regions are strongest among north Atlantic populations. The LG01 signal is particularly concentrated in the terminal branches leading to the Icelandic and Barents Sea branches. The LG02 signal is concentrated in the Icelandic terminal branch and the parent branches of the east Atlantic / north European. This region also contains a low-differentiation region inside it, suggesting it may be composed of two contiguous structural variants, as the LG01 region is known to be [85]. The LG07 signal is concentrated in nearly the same branches as the LG02 signal, and also in the Faroe plateau terminal branch. In contrast, the highly differentiated region in LG12 is concentrated among more downstream branches of the east Atlantic / north European populations, including the Celtic Sea terminal branch (Figure 6). Notably, none of the highly differentiated regions appears to show strong signs of high differentiation in the west Atlantic / North American populations.

## Discussion

We have developed a method for detecting positive selection when working with species with complex histories. The method is fast—it only took 486 seconds to run the bovine scan (including 512,358 SNPs and 36 populations) on a MacBook Air with a 1.8 GHz Intel Core i5 processor and 8 Gb of memory. Assuming a null model of genetic drift based on a multivariate Normal distribution, the *S_B_* statistic is chi-squared distributed with 1 degree of freedom. This is accurate as long as the graph topology is accurate and the branches in the graph do not contain high amounts of drift. In an admixture graph with *K* branches, there are *K* possible versions of the *S_B_* statistic. If the differences in allele frequencies at a SNP can be explained by an allele frequency shift that occurred along branch *k*, then *S_B_* (*k*) will be large, and a P-value based on the null drift model can be calculated from it. By design, branches whose parent are the root node and branches that have the same descendant nodes have the same *S_B_* scores, so selective events on these branches are not distinguishable from each other under this scheme.

The *S_B_* statistic is most accurate when a large number of individuals have been sampled from each population. If this is not the case, then one can average the scores over windows of SNPs to obtain power from correlated allele frequency shifts in a region (for example, as in ref. [26]), at the expense of losing spatial resolution across the genome due to larger test regions, as we did here in the bovine dataset example. The statistic, however, does not account for the structure of linkage disequilibrium within or between windows.

We have found the method performs best when there are many leaves in a graph because it uses a population-averaged allele frequency to estimate the ancestral allele frequency in the graph. We therefore recommend using this method when working with more than a few populations at a time, to make this estimate as accurate as possible. A possible future improvement of the method could be the incorporation of a model-based ancestral allele frequency estimation scheme, to address this issue. For example, to more appropriately account for the population covariance structure, we could also use the unbiased linear estimate of *e* with minimum variance, *ê* = (**1***^T^***F**^−1^**p**)/(**1***^T^***F**^−1^**1**) [14]. However, we found this estimate generates an excess of significant P-values when working with allele frequencies at the boundaries of fixation and extinction, due to poor modelling of frequency dynamics by the multivariate normal distribution, and that this effect is ameliorated when using the simpler population-averaged allele frequency.

Another critical issue is that the more branches one tests, the more of a multiple-testing burden there will be when defining significance cutoffs. In a way, our method improves on previous approaches to this problem, because—given an inferred admixture graph—one does not need to perform a test for all possible triplets or pairs of populations, as one would need to do when applying the PBS statistic or pairwise *F_ST_* methods, respectively. Instead, our method performs one test per branch. For example, if the graph is a rooted tree with *m* leaves and no admixture, the number of branches will be equal to 2*m* – 2, while the number of possible triplets will be equal to 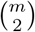, and the number of pairs will be equal to 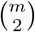, both of which grow much faster with larger m than does 2*m* – 2.

While the SB statistic is fast and easy to compute, it is not as principled as other approaches for multi-population selection that rely on explicit models of positive selection (e.g. [86]). This means that it only detects significant deviations from a neutral null model and does not provide likelihoods or posterior probabilities supporting specific selection models. We recommend that, once a locus with high *S_B_* has been detected in a particular branch of a graph, biologists should perform further work to disentangle exactly what type of phenomenon would cause this value to be so high and test among competing selection hypotheses.

Among the genes that emerge when applying our method to human data, we found several known candidates, like LCT/MCM6, SLC45A2, SLC24A5, POU2F3, OCA2/HERC2 and BNC2. We also found several new candidate regions, containing genes involved in the immune response, like the TARBP1 and NFAM1 genes in East Asians. Additionally, we found new candidate regions in Native Americans, like GSK3B and the protamine gene cluster.

Analysis of the bovine dataset yielded numerous regions that may be implicated in the breeding process. One of the strongest candidate regions contains genes involved in musculoskeletal morphology, including CEP120 and PRDM6, and *GRoSS* narrows this signal down to the branch leading to the Maremmana breed. This is an Italian beef cattle breed that inhabits the Maremma region in Central Italy, and has evolved a massive body structure well adapted to draft use in the marshy land that characterizes the region [87]. Interestingly, when comparing muscle samples between Maremmana and the closely-related Chianina breed (CH), gene ontology categories related to muscle structural proteins and regulation of muscle contraction have been reported to be enriched for differentially expressed genes. Additionally, the Maremmana is enriched for over-expressed genes related to hypertrophic cardiomyopathy pathways [87].

Another strong candidate region is the protocadherin gene cluster, associated with neuronal functions in humans and mice [75–77], and shown to be under positive selection in domesticated cats and foxes [78, 79]. *GRoSS* identifies this region as under selection in the Romanian Grey breed terminal branch. Given that this breed is popularly known to be very docile, it is plausible that this gene cluster might have been a target for selection on behavior during the recent breeding process.

Additionally, *GRoSS* detects a very large 4.4-Mb region as a selection candidate in the Holstein breed, currently the world’s highest-production dairy animal. This region overlaps several candidate genes earlier identified to be under selection in Holstein using other methods (see [72] for an extensive review). These genes are related to several traits usually targeted by breeding practices, such as behaviour, muscle development and milk yield.

Our method also recovered previously reported regions of high differentiation among Atlantic codfish populations and served to pinpoint where in the history of this species the inversions may have arisen, or at least where they have most strongly undergone the process of differentiation between haplotypes. The largest of these regions is in LG01 and is composed of two adjacent inversions covering 17.4 Mb [85], which suppress recombination in heterozygous individuals and promote differentiation between haplotypes. The inversions effectively lock together a super-gene of alleles at multiple loci [85]. Two behavioral ecotypes—a deep-sea frontal (migratory) ecotype and a shallow-water coastal (stationary) ecotype—have been associated with inversion alleles in the region [88–90]. Several putative candidate selected genes are located within the LG01 inversions [85, 91, 92] that may be of adaptive value for deep sea as well as long-distance migration.

Similarly, the other large inversions observed on linkage groups LG02, LG07 and LG12 (5, 9.5, and 13 Mb respectively) also suppress recombination [93, 94]. Allele frequency differences observed between individuals living offshore and inshore environments are suggestive of ecological adaptation driving differentiation in these regions [93–95]. Previously, a pairwise *F_ST_* outlier analysis of populations in the north (Greenland, Iceland, and Barents Sea localities combined) vs. populations in the south (Faroe Islands, North Sea, and Celtic Sea combined) showed clear evidence of selection in these regions [48]. However, in comparisons of West (Sable Bank, Western Bank, Trinity Bay, and Southern Grand Banks combined) with either North or South localities only some of these regions displayed signatures of high differentiation [48], indicating these inversions had different spatiotemporal origins. By modeling all these populations together in a single framework, our method provides a way to more rigorously determine in which parts of the graph these inversions may have originated (Figure 6), and suggests they were largely restricted to East Atlantic populations.

In conclusion, *GRoSS* is a freely-available, fast and intuitive approach to testing for positive selection when the populations under study are related via a history of multiple population splits and admixture events. It can identify signals of adaptation in a species by accounting for the complexity of this history, while also providing a readily interpretable score. This method will help evolutionary biologists and ecologists pinpoint when and where adaptive events occurred in the past, facilitating the study of natural selection and its biological consequences.

## Acknowledgments

We thank Jeremy Berg, Anders Albrechtsen and Kathleen Lotterhos for helpful advice and discussions. FR thanks the Natural History Museum of Denmark, the Danish National Research Foundation and the Lundbeck Foundation for their support. EÁ and KH were supported by a grant from Svala Árnadóttir’s private funds, by a grant from the University of Iceland Research Fund, by institutional funds from R.C. Lewontin, and by a grant from the Icelandic Research Fund (no. 185151–051). R.R.F. thanks the Danish National Research Foundation for its support of the Center for Macroecology, Evolution, and Climate (grant DNRF96). The authors declare no competing interests.

## Supplementary Figures

**Figure S1:**
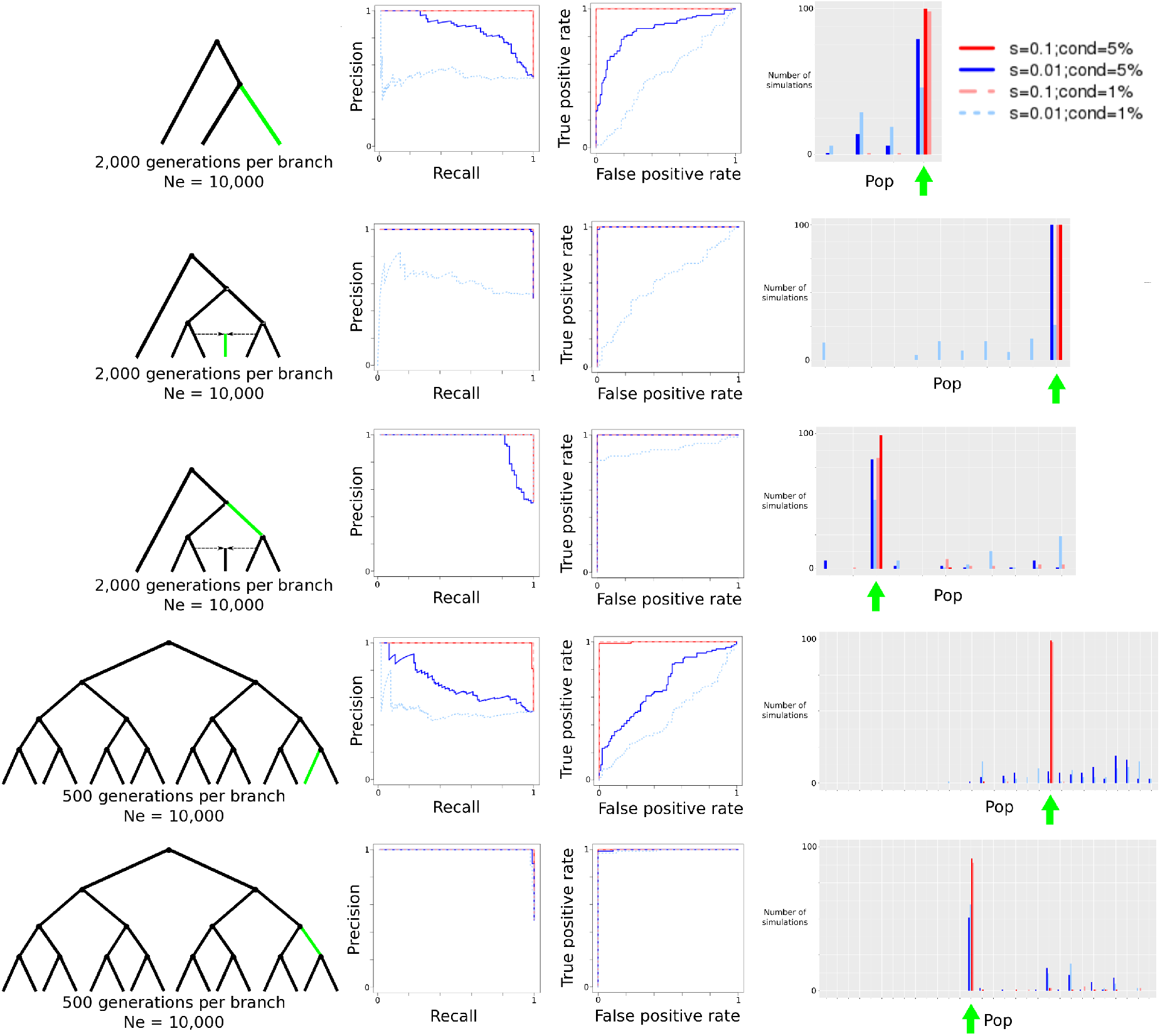
Evaluation of *GRoSS* performance using simulations in *SLiM 2*, with 50 diploid invididuals per population panel. We simulated different selective sweeps under strong (s=0.1) and intermediate (s=0.01) selection coefficients for a 3-population tree, a 6-population graph with a 50%/50% admixture event and a 16-population tree. We then produced precision-recall and ROC curves comparing simulations under selection to simulations under neutrality. We also obtained the maximum branch score within 100kb of the selected site, and computed the number of simulations (out of 100) in which the branch of this score corresponded to the true selected branch. “*cond* = 5%”: Simulations conditional on the beneficial mutation reaching 5% frequency or more. “*cond* = 1%”: Simulations conditional on the beneficial mutation reaching 1% frequency or more. “Pop”: population branch.

**Figure S2:**
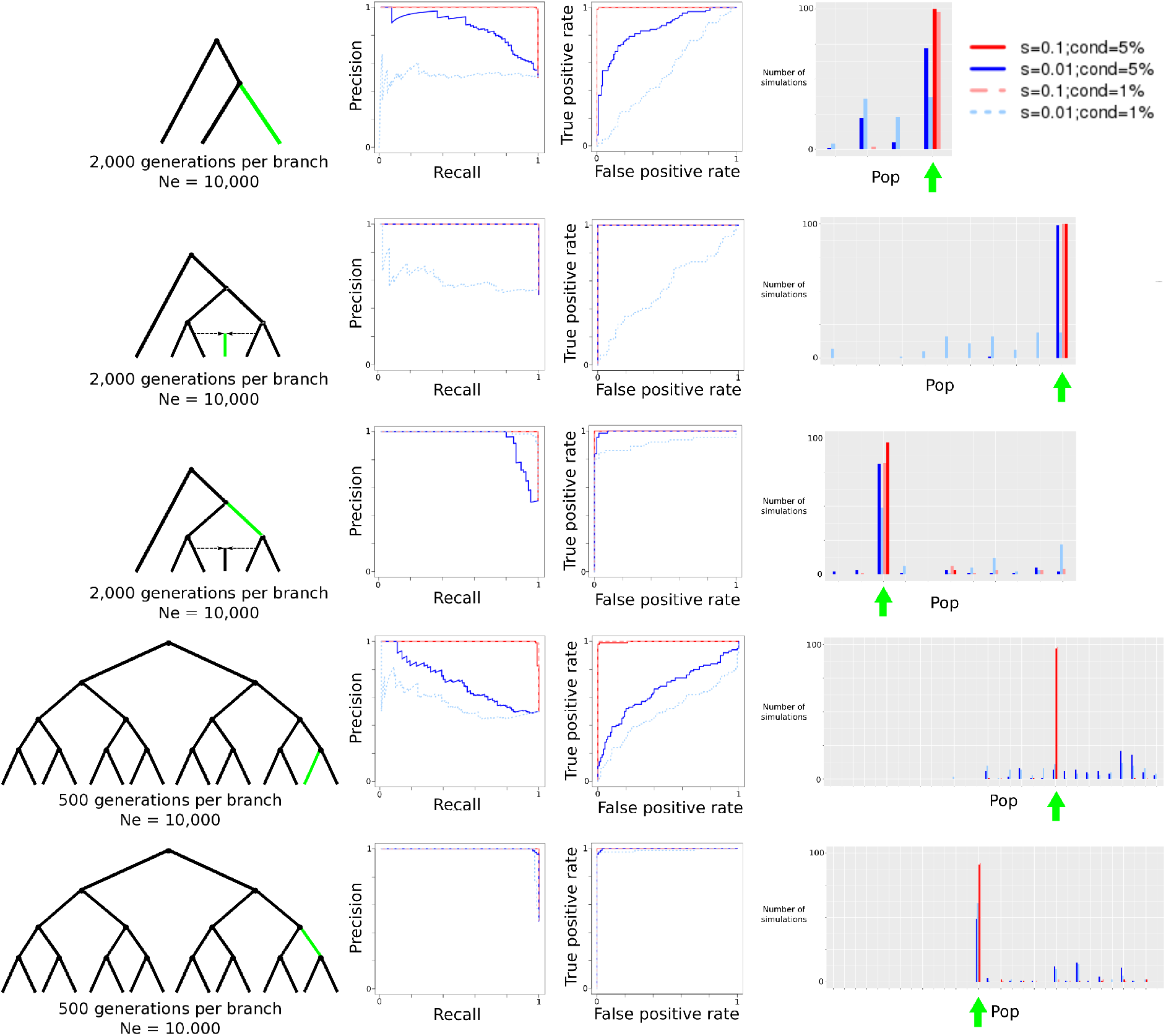
Evaluation of *GRoSS* performance using simulations in *SLiM* 2, with 25 diploid individuals per population panel. We simulated different selective sweeps under strong (s=0.1) and intermediate (s=0.01) selection coefficients for a 3-population tree, a 6-population graph with a 50%/50% admixture event and a 16-population tree. We then produced precision-recall and ROC curves comparing simulations under selection to simulations under neutrality. We also obtained the maximum branch score within 100kb of the selected site, and computed the number of simulations (out of 100) in which the branch of this score corresponded to the true selected branch. “*cond* = 5%”: Simulations conditional on the beneficial mutation reaching 5% frequency or more. “*cond* = 1%”: Simulations conditional on the beneficial mutation reaching 1% frequency or more. “Pop”: population branch.

**Figure S3:**
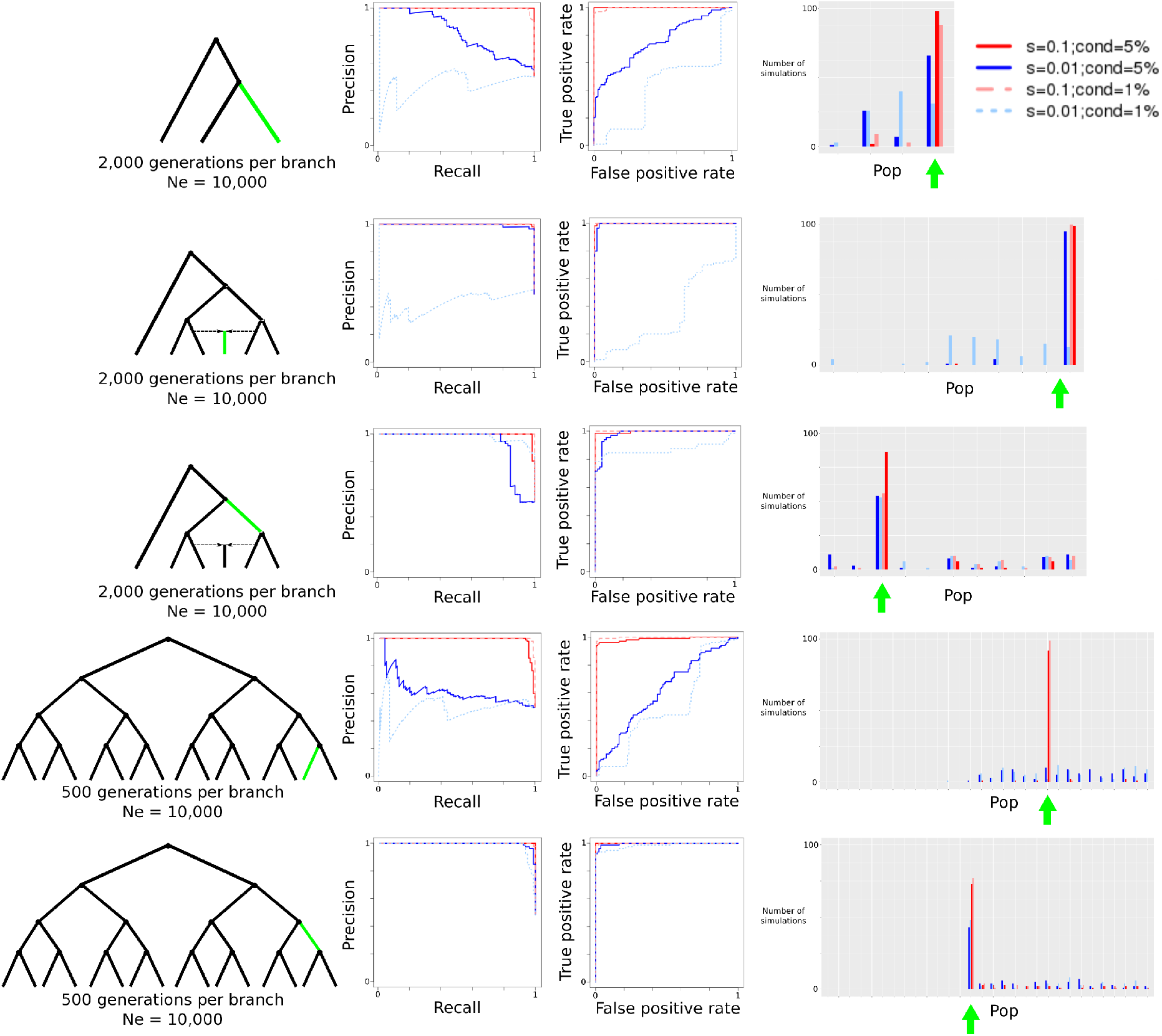
Evaluation of *GRoSS* performance using simulations in *SLiM* 2, with 4 diploid individuals per population panel. We simulated different selective sweeps under strong (s=0.1) and intermediate (s=0.01) selection coefficients for a 3-population tree, a 6-population graph with a 50%/50% admixture event and a 16-population tree. We then produced precision-recall and ROC curves comparing simulations under selection to simulations under neutrality. We also obtained the maximum branch score within 100kb of the selected site, and computed the number of simulations (out of 100) in which the branch of this score corresponded to the true selected branch. “*cond* = 5%”: Simulations conditional on the beneficial mutation reaching 5% frequency or more. “*cond* = 1%”: Simulations conditional on the beneficial mutation reaching 1% frequency or more. “Pop”: population branch.

**Figure S4:**
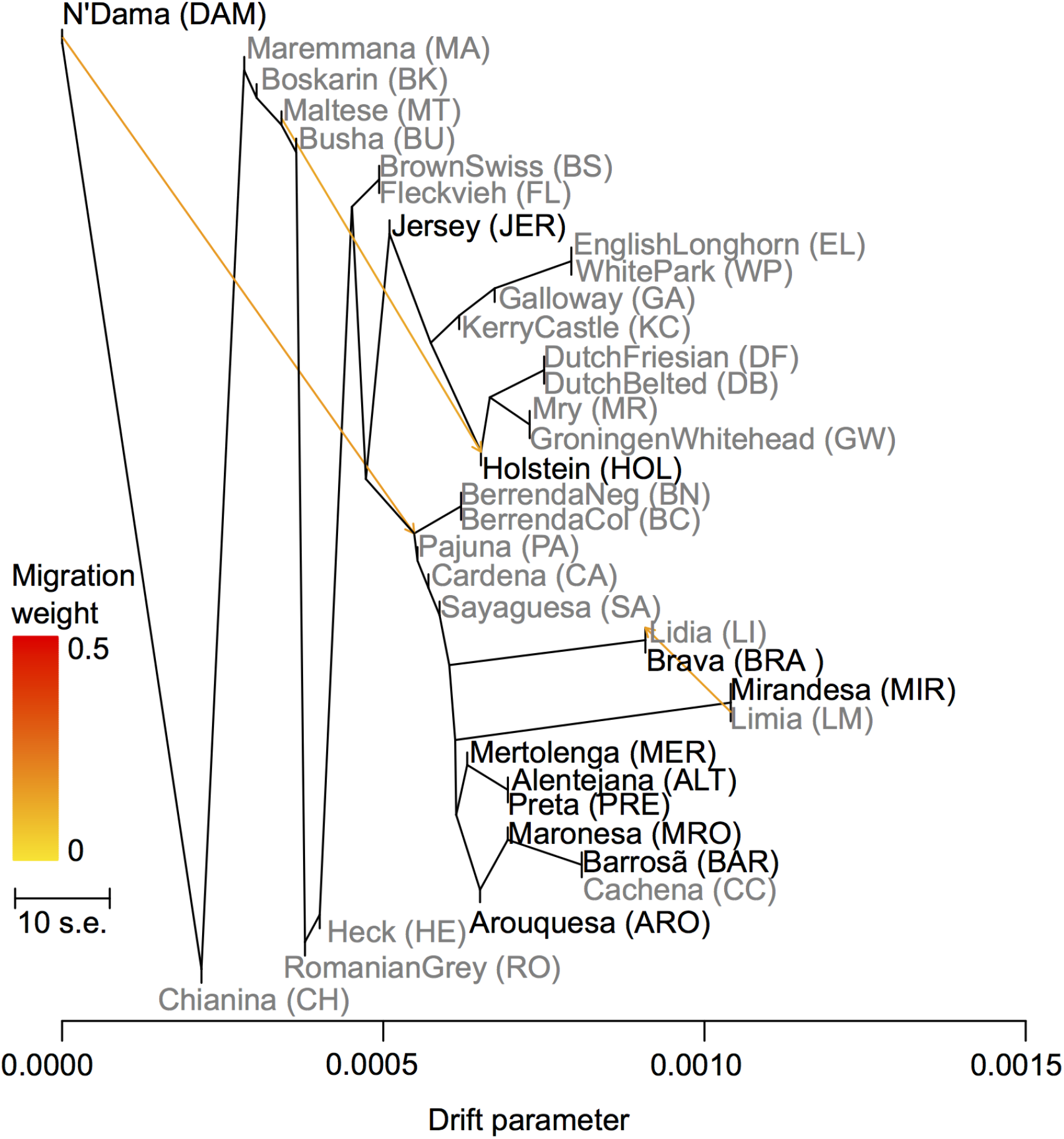
*TreeMix*-fitted maximum likelihood admixture graph with 3 admixture events, depicting the relationships between the taurine cattle breeds analyzed in this study (grey: Illumina BovineHD SNP data; black: whole genome data).

**Figure S5:**
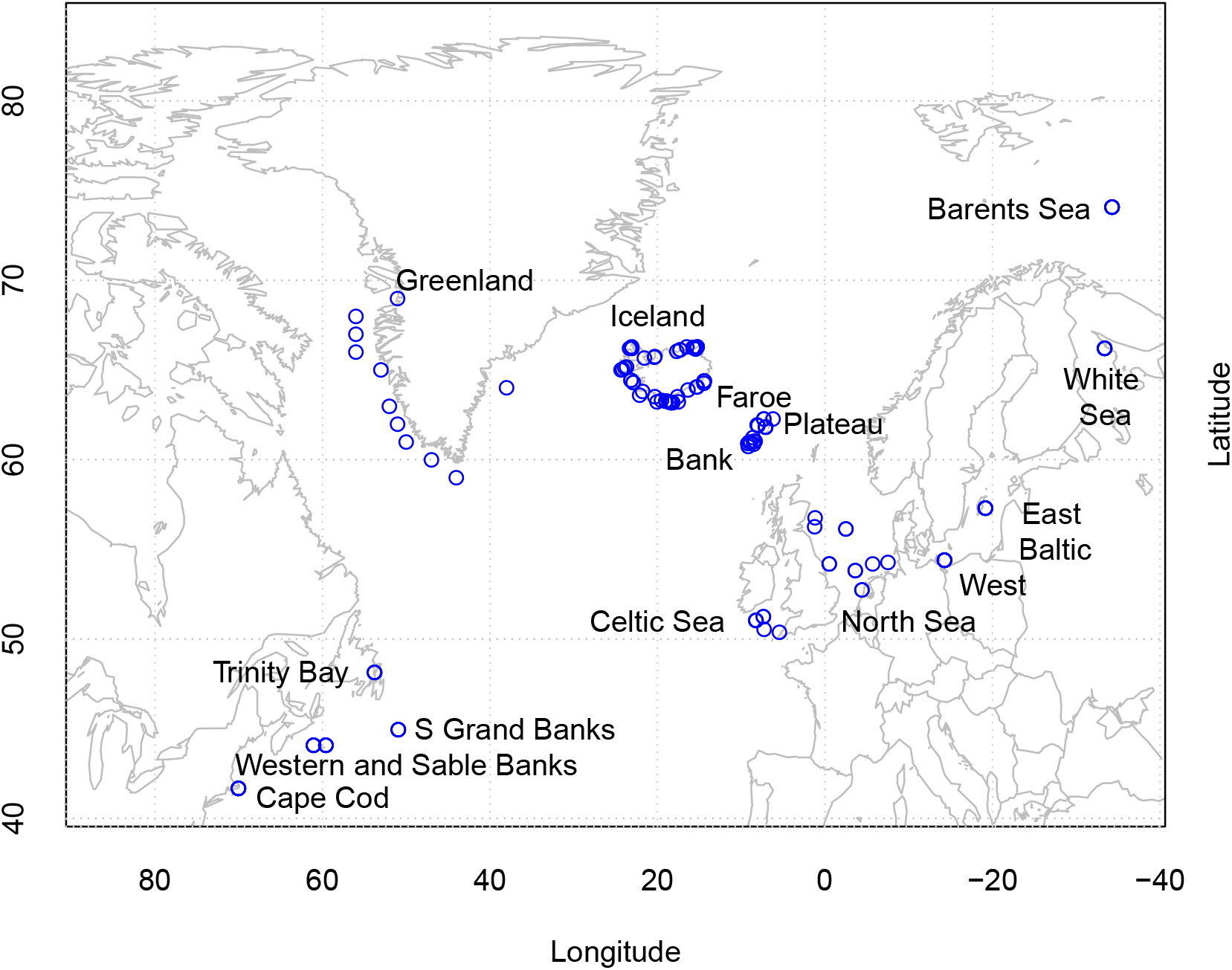
Sample localities of Atlantic cod samples on a map of the North Atlantic.

**Figure S6:**
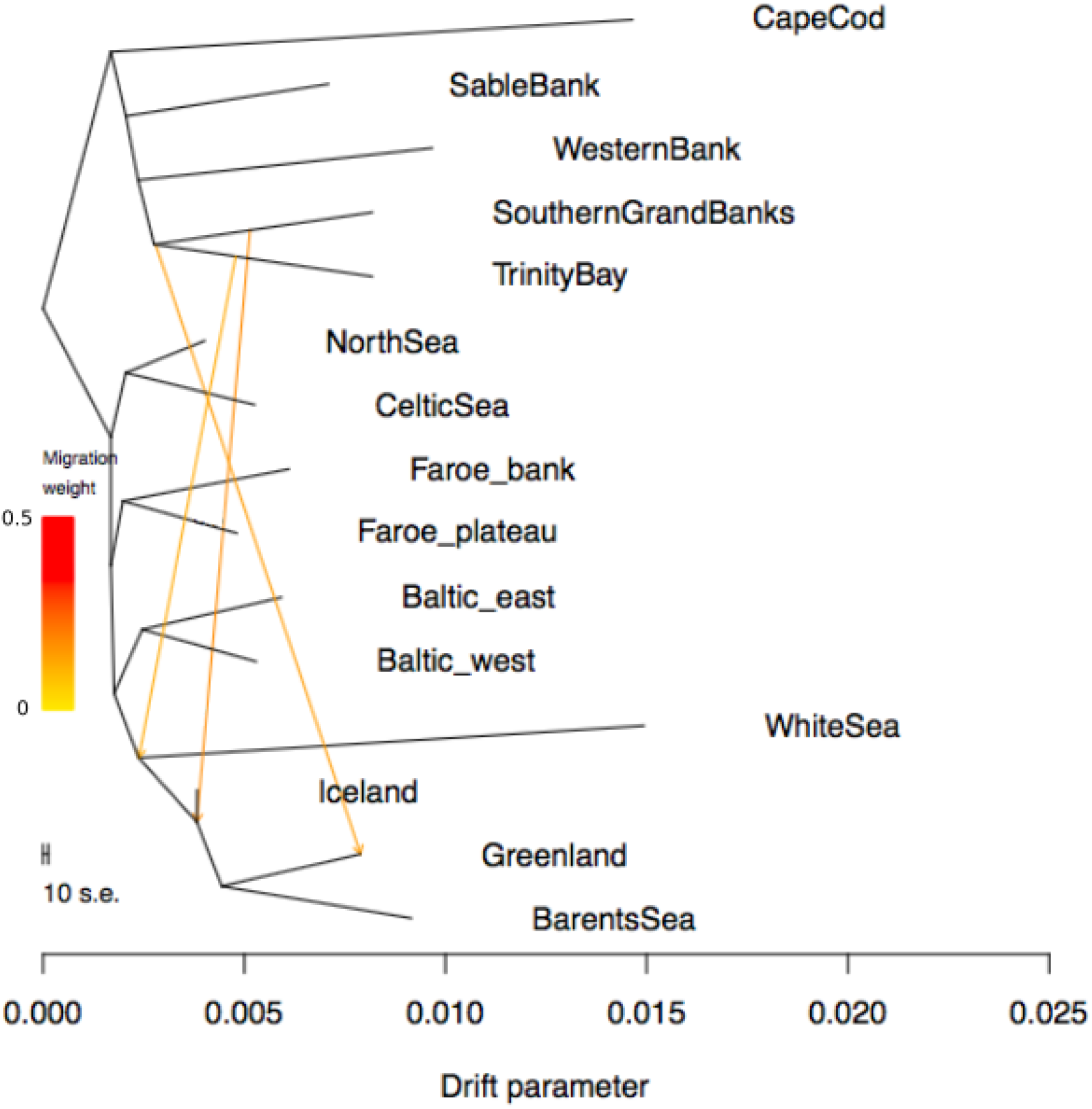
*Treemix*-fitted maximum likelihood admixture graph with 3 admixture events, depicting the relationships between the Atlantic codfish populations.

**Figure S7:**
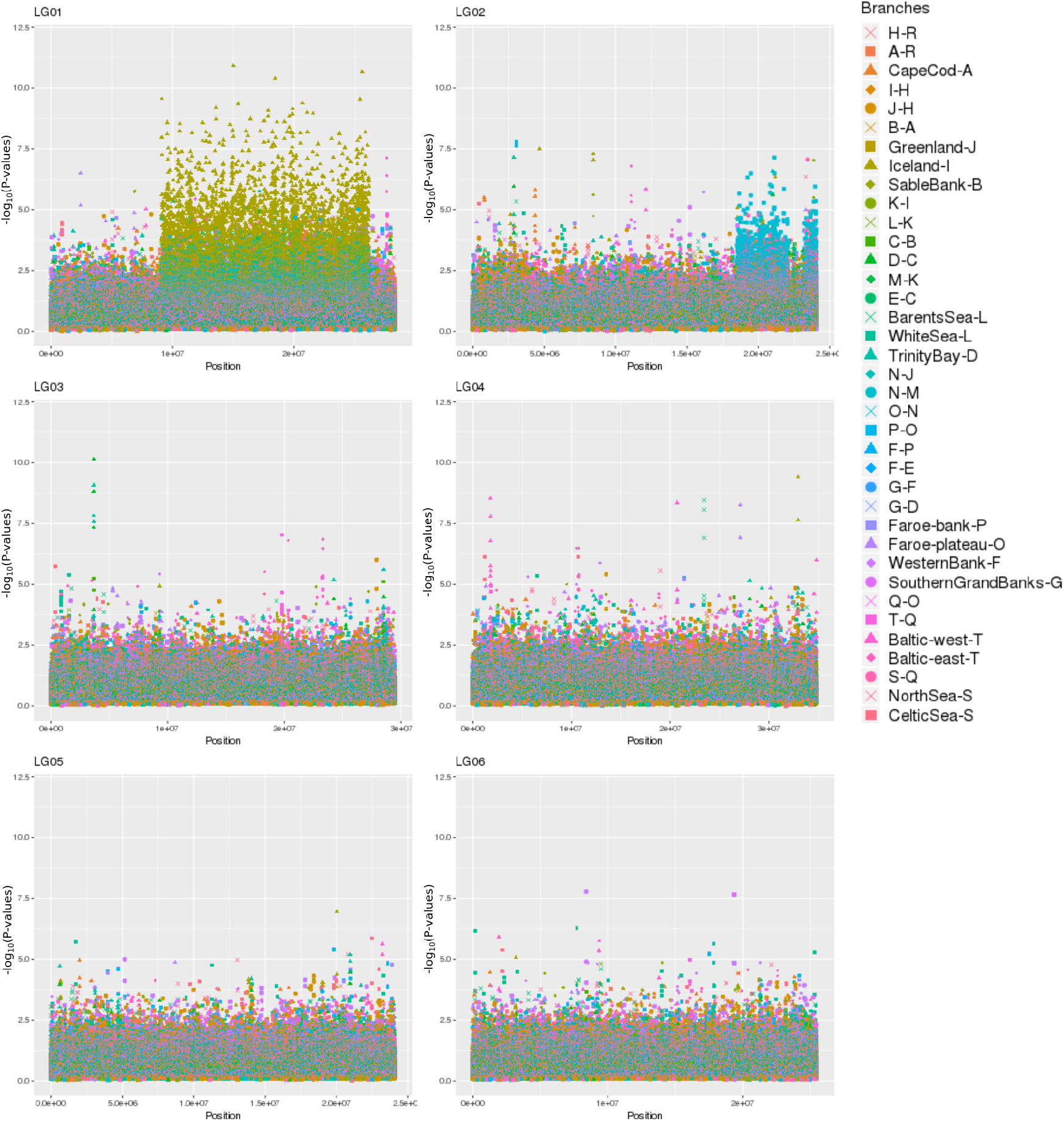
*S_B_* scores for LG01 - LG06 in the Codfish data.

**Figure S8:**
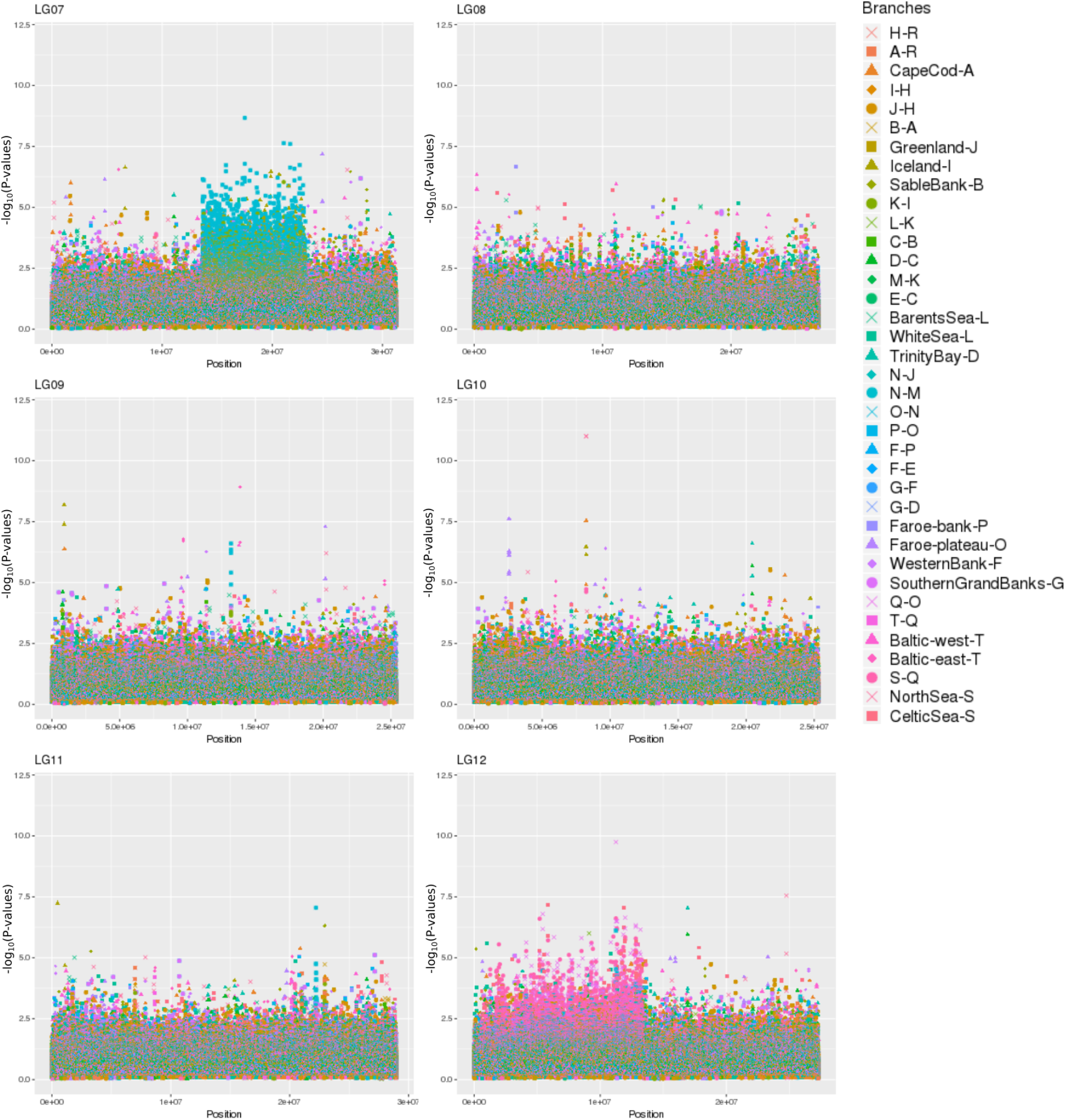
*S_B_* scores for LG07 - LG12 in the Codfish data.

**Figure S9:**
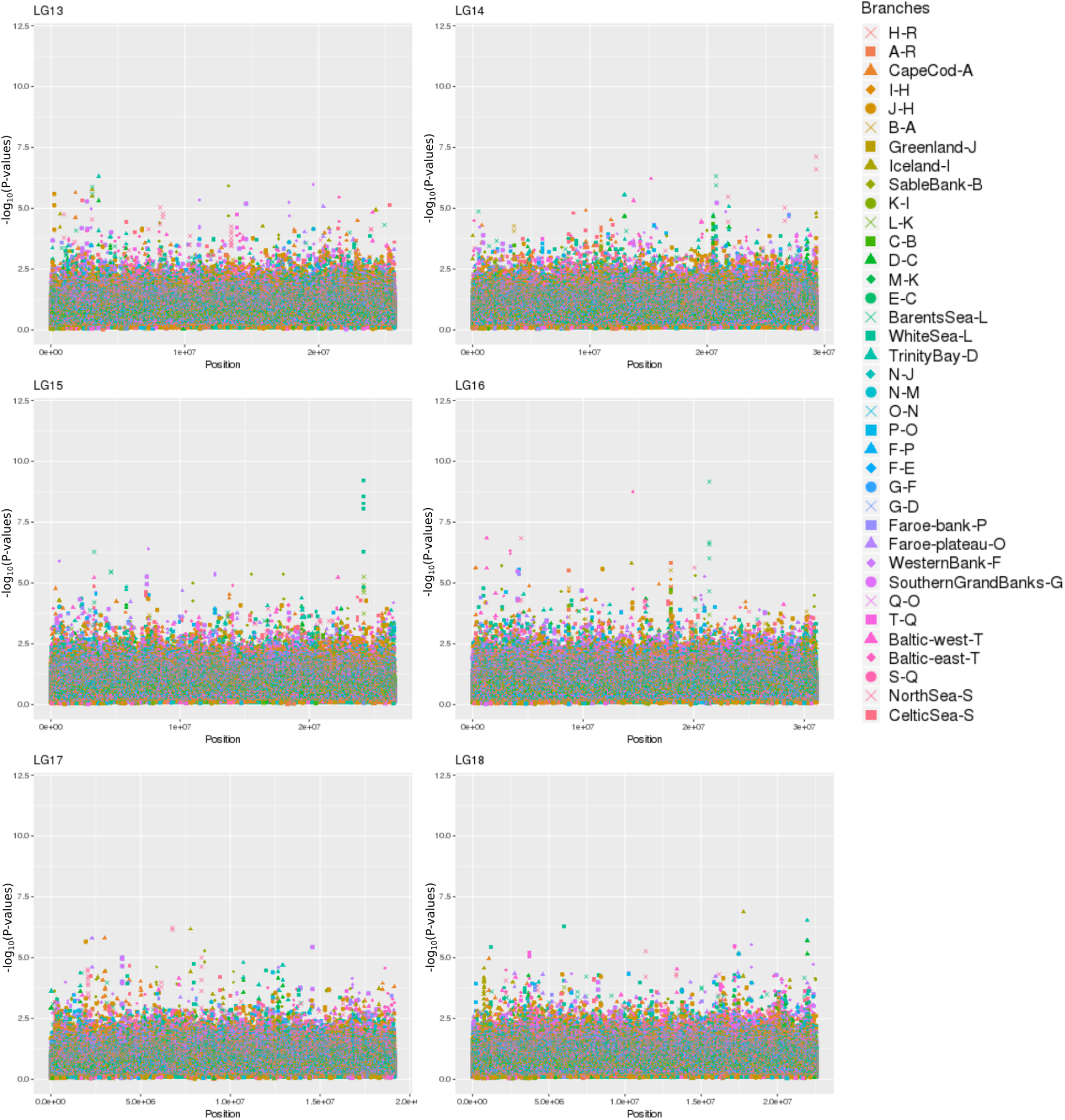
*S_B_* scores for LG13 - LG18 in the Codfish data.

**Figure S10:**
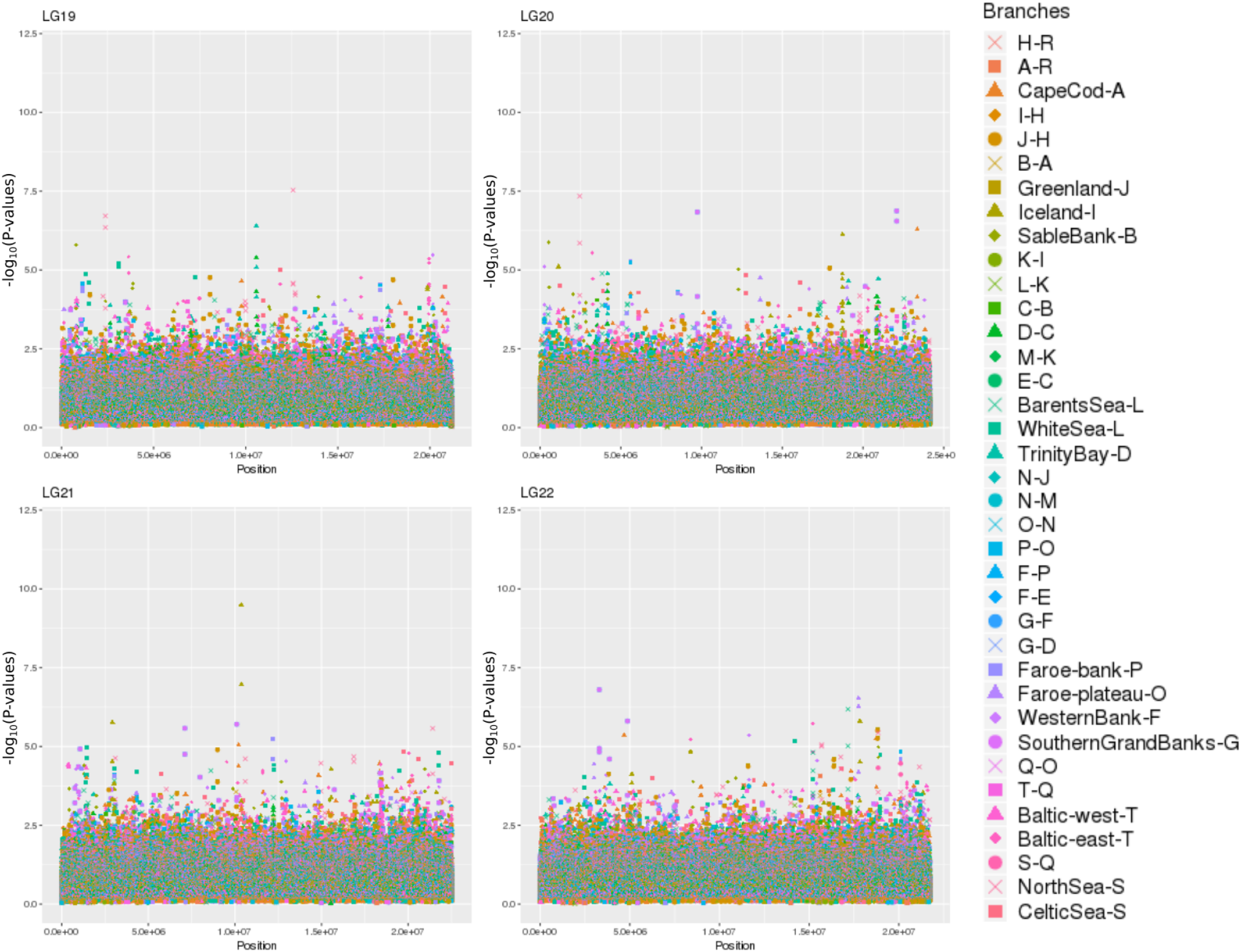
*S_B_* scores for LG19 - LG22 in the Codfish data.

## Supplementary Tables

**Table S1:**
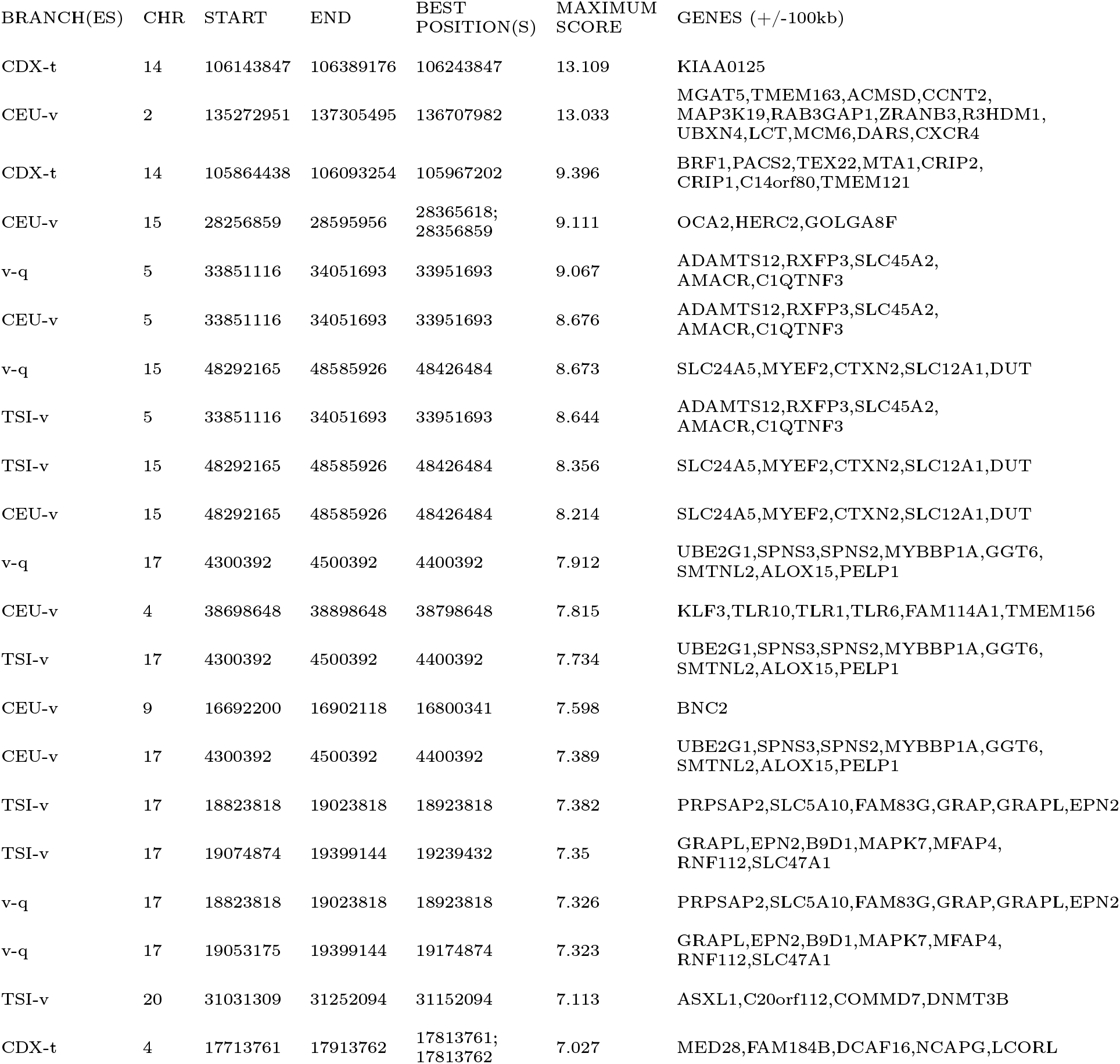
Top candidate regions from 1000 Genomes scan.

**Table S2:**
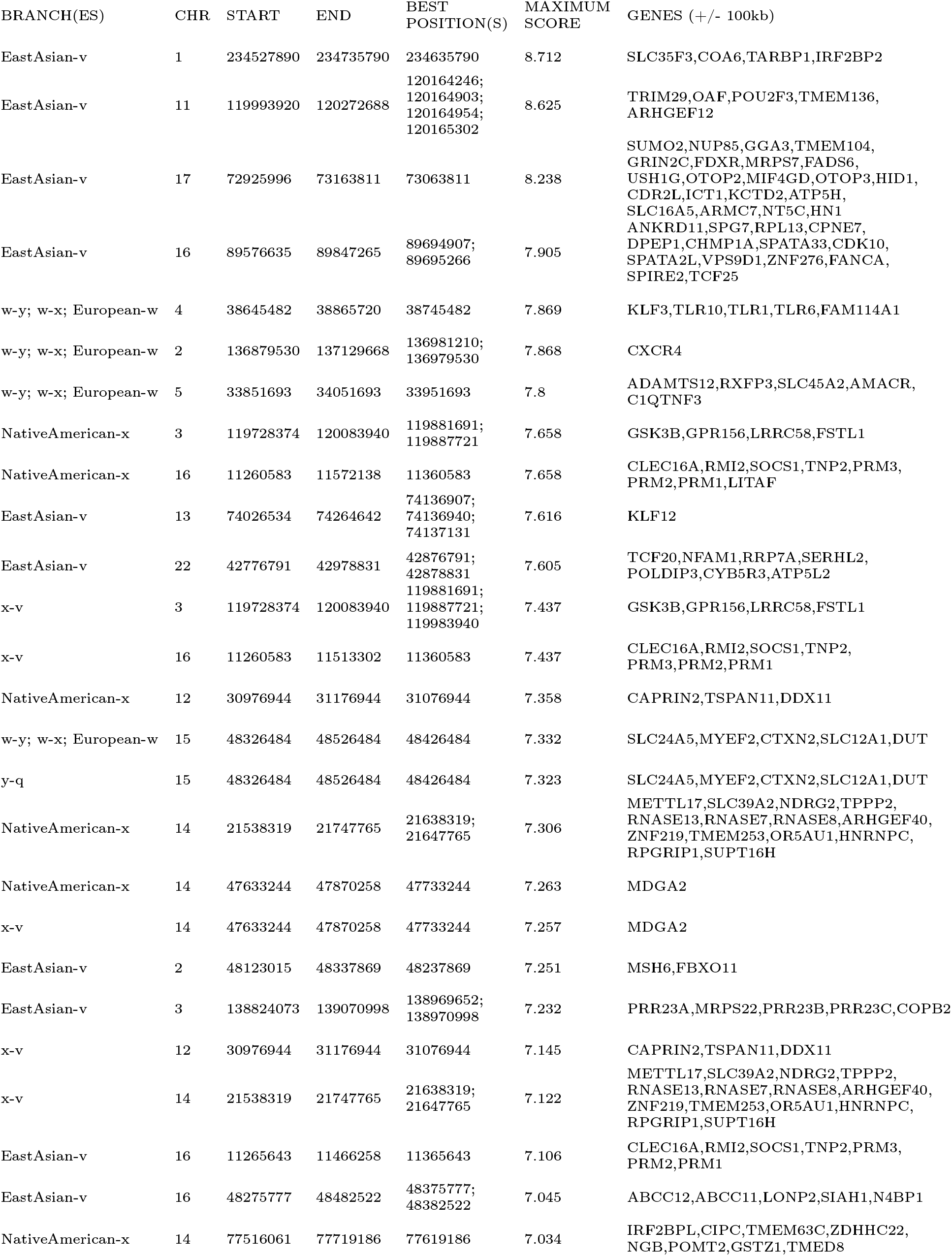
Top candidate regions from Human Origins scan.

**Table S3:** Country, sample sizes and data type for panels of cattle breeds analyzed in this study.

| Abbreviation | Breed name | Country or region | Sample size | Data type | Reference |
| --- | --- | --- | --- | --- | --- |
| ALT | Alentejana | Portugal | 6 | shotgun | da Fonseca et al. (submitted) |
| ARO | Arouquesa | Portugal | 6 | shotgun | da Fonseca et al. (submitted) |
| BAR | Barrosã | Portugal | 6 | shotgun | da Fonseca et al. (submitted) |
| BC | Berrenda en colorado | Spain | 3 | 777K chip | Upadhyay et al. 2017 |
| BK | Boskarin | Hungary | 4 | 777K chip | Shaheen et al. 2015 |
| BN | Berrenda en negro | Spain | 3 | 777K chip | Upadhyay et al. 2017 |
| BRA | Brava de Lide | Portugal | 6 | shotgun | da Fonseca et al. (submitted) |
| BS | Brown Swiss | Switzerland | 4 | 777K chip | Upadhyay et al. 2017 |
| BU | Busha | Balkan region | 6 | 777K chip | Upadhyay et al. 2017 |
| CA | Cachena | Portugal | 3 | 777K chip | Upadhyay et al. 2017 |
| CCIBR | Cardena | Spain | 5 | 777K chip | Upadhyay et al. 2017 |
| CH | Chianina | Italy | 3 | 777K chip | Upadhyay et al. 2017 |
| DAM | N'Dama | Africa | 10 | shotgun | Kim et al. 2017 |
| DB | Dutch Belted | The Netherlands | 2 | 777K chip | Upadhyay et al. 2017 |
| DF | Dutch Friesian | The Netherlands | 4 | 777K chip | Upadhyay et al. 2017 |
| EL | English Longhorn | England | 4 | 777K chip | Upadhyay et al. 2017 |
| FL | Fleckvieh | Switzerland | 4 | 777K chip | Upadhyay et al. 2017 |
| GA | Galloway | Scotland | 5 | 777K chip | Upadhyay et al. 2017 |
| GW | Groningen Whiteheaded | The Netherlands | 5 | 777K chip | Upadhyay et al. 2017 |
| HE | Heck | Germany | 5 | 777K chip | Upadhyay et al. 2017 |
| HOL | Holstein | The Netherlands | 10 | shotgun | Kim et al. 2017 |
| JER | Jersey | Jersey Island | 9 | shotgun | Kim et al. 2017 |
| KC | Kerry Cattle | Ireland | 4 | 777K chip | Upadhyay et al. 2017 |
| LI | Lidia | Spain | 3 | 777K chip | Upadhyay et al. 2017 |
| LM | Limia | Spain | 4 | 777K chip | Upadhyay et al. 2017 |
| MA | Maremmana | Italy | 5 | 777K chip | Upadhyay et al. 2017 |
| MER | Mertolenga | Portugal | 6 | shotgun | da Fonseca et al. (submitted) |
| MIR | Mirandesa | Portugal | 6 | shotgun | da Fonseca et al. (submitted) |
| MR | Meuse-Rhine-Yssel | The Netherlands | 4 | 777K chip | Upadhyay et al. 2017 |
| MRO | Maronesa | Portugal | 6 | shotgun | da Fonseca et al. (submitted) |
| MT | Maltese | Malta | 4 | 777K chip | Upadhyay et al. 2017 |
| PA | Pajuna | Spain | 6 | 777K chip | Upadhyay et al. 2017 |
| PRE | Preta | Portugal | 6 | shotgun | this study |
| RO | Romanian grey | Romania | 4 | 777K chip | Upadhyay et al. 2017 |
| SA | Sayaguesa | Spain | 5 | 777K chip | Upadhyay et al. 2017 |
| WP | White Park | England | 3 | 777K chip | Upadhyay et al. 2017 |

**Table S4:**
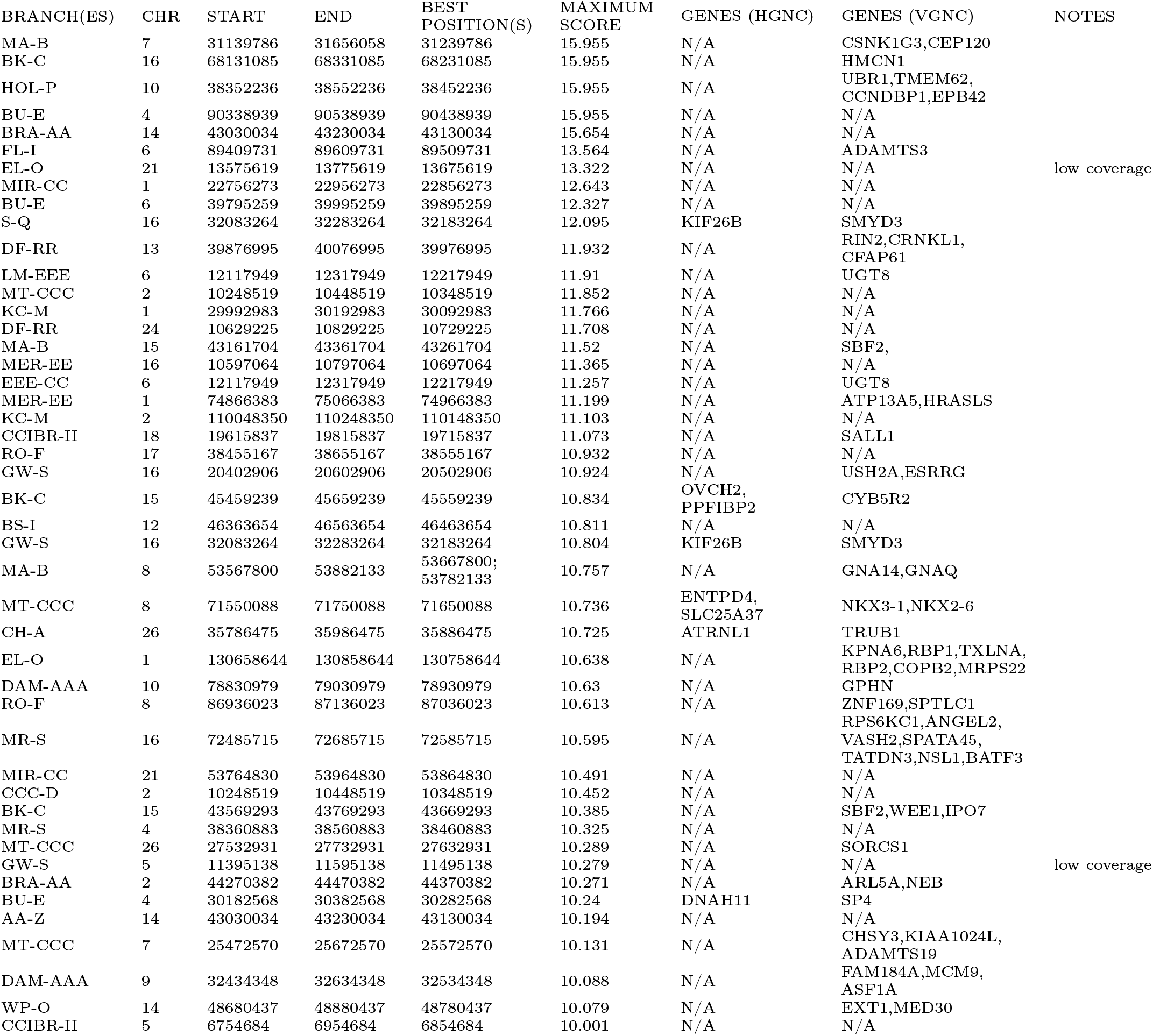
Top candidate regions from bovine selection scan, computing 1 score per SNP. Gene annotations were extracted from the Human Gene Nomenclature Committee (HGNC) and the Vertebrate Gene Nomenclature Committee (VGNC). We labeled windows with particularly low coverage in the target population, as the signal of selection may be inflated in those windows for that reason.

**Table S5:**
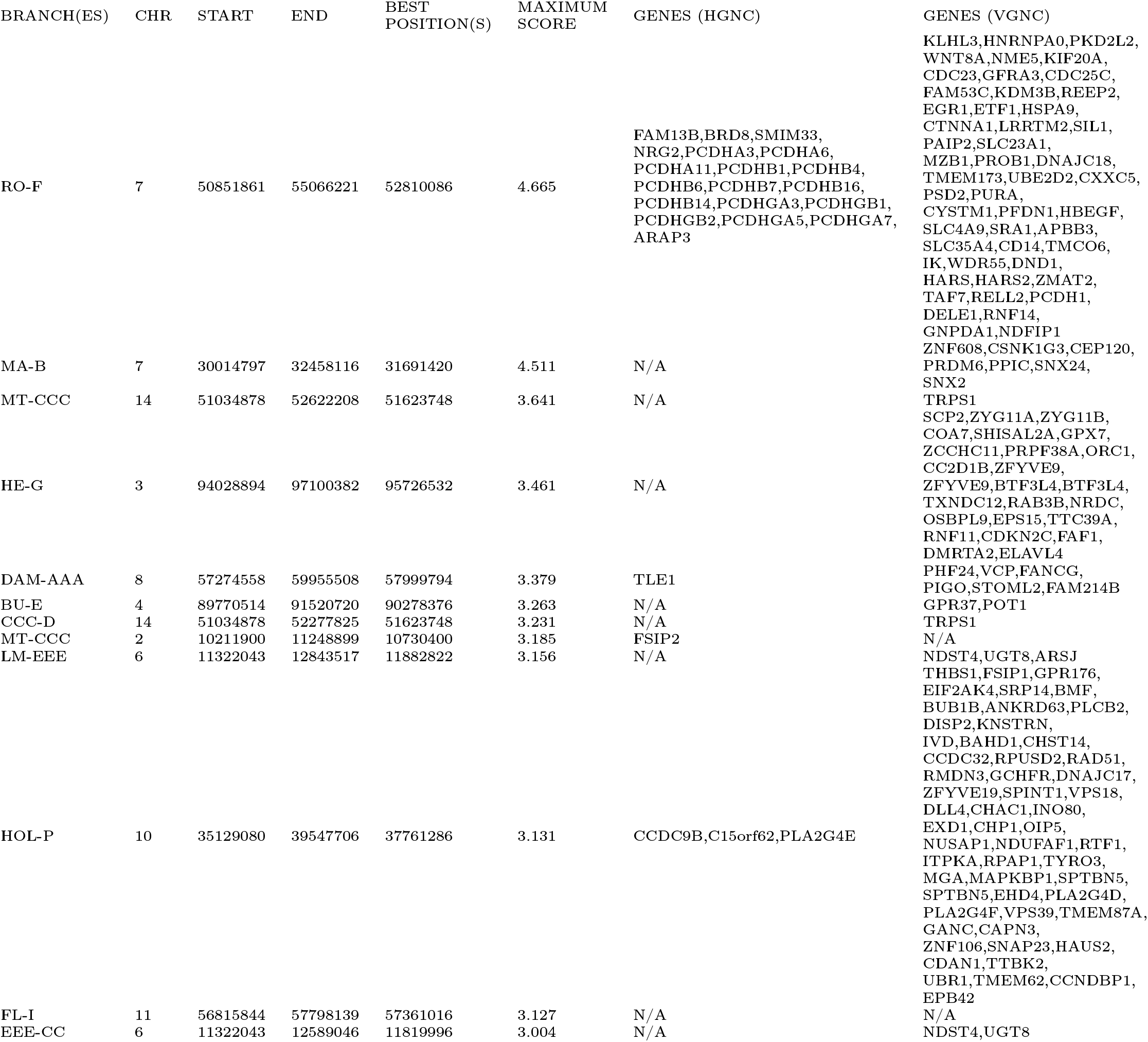
Top candidate regions from bovine selection scan, computing score in windows of 10 SNPs with a step size of 1 SNP. Gene annotations were extracted from the Human Gene Nomenclature Committee (HGNC) and the Vertebrate Gene Nomenclature Committee (VGNC).

**Table S6:** Area of sampling and sample sizes of Atlantic cod population panels analyzed in this study (see Figure S5).

| Area | Population | Abbreviation | Sample size |
| --- | --- | --- | --- |
| US | Cape Cod | Cco | 6 |
| Nova Scotia | Western Bank | Web | 5 |
| Nova Scotia | Sable Bank | Sab | 5 |
| Newfoundland | Trinity Bay | Tri | 4 |
| Newfoundland | Southern Grand Banks | Sgb | 4 |
| Greenland | Greenland | Gre | 11 |
| Iceland | Iceland | Ice | 61 |
| Norway | Barents Sea | Bar | 8 |
| Faroes | Faroe Plateau | FarP | 8 |
| Faroes | Faroe Bank | FarB | 5 |
| Celtic Sea | Celtic Sea | Cel | 8 |
| North Sea | North Sea | Nse | 12 |
| Baltic Sea | Baltic West | BalW | 8 |
| Baltic Sea | Baltic East | BalE | 8 |
| Russia | White Sea | Whi | 8 |

**Table S7:**
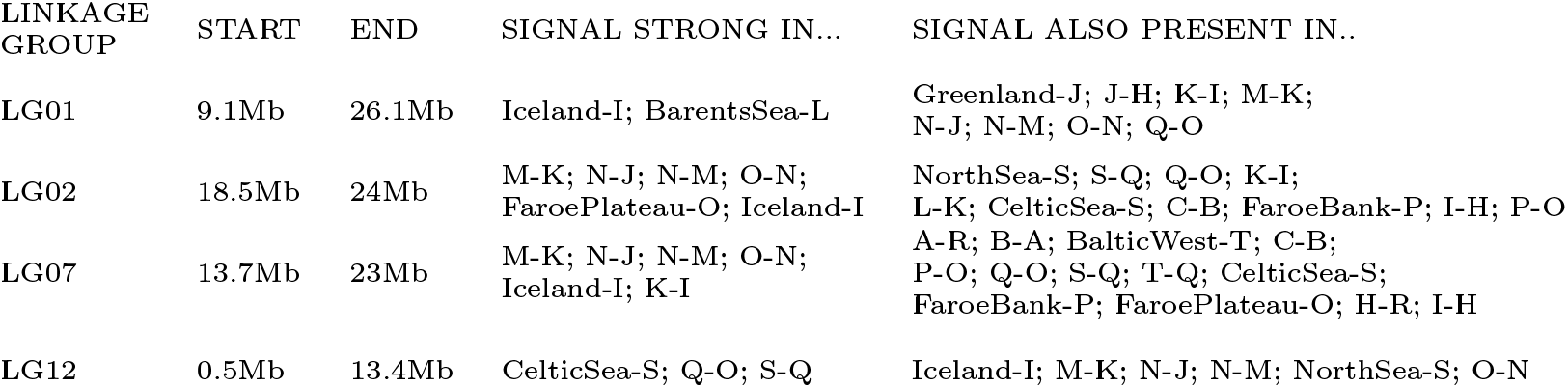
Long high-differentiation regions in the Codfish data. Branches with corresponding scores with –*log*_10_(*P*) > 5 for at least 1 SNP inside the region were placed into the column ‘SIGNAL STRONG IN…’.

| LINKAGE GROUP | START | END | SIGNAL STRONG IN... | SIGNAL ALSO PRESENT IN.. |
| --- | --- | --- | --- | --- |
| LG01 | 9.1Mb | 26.1Mb | Iceland-I; BarentsSea-L | Greenland-J; J-H; K-I; M-K; N-J; N-M; O-N; Q-O |
| LG02 | 18.5Mb | 24Mb | M-K; N-J; N-M; O-N; FaroePlateau-O; Iceland-I | NorthSea-S; S-Q; Q-O; K-I; L-K; CelticSea-S; C-B; FaroeBank-P; I-H; P-O |
| LG07 | 13.7Mb | 23Mb | M-K; N-J; N-M; O-N; Iceland-I; K-I | A-R; B-A; BalticWest-T; C-B; P-O; Q-O; S-Q; T-Q; CelticSea-S; FaroeBank-P; FaroePlateau-O; H-R; I-H |
| LG12 | 0.5Mb | 13.4Mb | CelticSea-S; Q-O; S-Q | Iceland-I; M-K; N-J; N-M; NorthSea-S; O-N |

## References

1. Voight, B. F., Kudaravalli, S., Wen, X. & Pritchard, J. K. A map of recent positive selection in the human genome. PLos biology 4, e72 (2006).

2. Sabeti, P. C. et al. Genome-wide detection and characterization of positive selection in human populations. Nature 449, 913 (2007).

3. Nielsen, R. et al. Genomic scans for selective sweeps using SNP data. Genome research 15, 1566–1575 (2005).

4. Huber, C. D., DeGiorgio, M., Hellmann, I. & Nielsen, R. Detecting recent selective sweeps while controlling for mutation rate and background selection. Molecular ecology 25, 142–156 (2016).

5. Shriver, M. D. et al. The genomic distribution of population substructure in four populations using 8,525 autosomal SNPs. Human genomics 1, 274 (2004).

6. Yi, X. et al. Sequencing of 50 human exomes reveals adaptation to high altitude. Science 329, 75–78 (2010).

7. Wright, S. The genetical structure of populations. Annals of Human Genetics 15, 323–354 (1949).

8. Weir, B. S. & Cockerham, C. C. Estimating F-statistics for the analysis of population structure. evolution 38, 1358–1370 (1984).

9. Lewontin, R. & Krakauer, J. Distribution of gene frequency as a test of the theory of the selective neutrality of polymorphisms. Genetics 74, 175–195 (1973).

10. Weir, B. S., Cardon, L. R., Anderson, A. D., Nielsen, D. M. & Hill, W. G. Measures of human population structure show heterogeneity among genomic regions. Genome research 15, 1468–1476 (2005).

11. Chen, H., Patterson, N. & Reich, D. Population differentiation as a test for selective sweeps. Genome research 20, 393–402 (2010).

12. Racimo, F. Testing for ancient selection using cross-population allele frequency differentiation. Genetics 202, 733–750 (2016).

13. Cheng, X., Xu, C. & DeGiorgio, M. Fast and robust detection of ancestral selective sweeps. Molecular ecology 26, 6871–6891 (2017).

14. Bonhomme, M. et al. Detecting selection in population trees: the Lewontin and Krakauer test extended. Genetics 186, 241–262 (2010).

15. Fariello, M. I., Boitard, S., Naya, H., SanCristobal, M. & Servin, B. Detecting signatures of selection through haplotype differentiation among hierarchically structured populations. Genetics 193, 929–941 (2013).

16. Librado, P. & Orlando, L. Detecting signatures of positive selection along defined branches of a population tree using LSD. Molecular biology and evolution 35, 1520–1535 (2018).

17. Coop, G., Witonsky, D., Di Rienzo, A. & Pritchard, J. K. Using environmental correlations to identify loci underlying local adaptation. Genetics 185, 1411–1423 (2010).

18. Günther, T. & Coop, G. Robust identification of local adaptation from allele frequencies. Genetics 195, 205–220 (2013).

19. Guillot, G., Vitalis, R., le Rouzic, A. & Gautier, M. Detecting correlation between allele frequencies and environmental variables as a signature of selection. A fast computational approach for genome-wide studies. Spatial Statistics 8, 145–155 (2014).

20. Villemereuil, P. & Gaggiotti, O. E. A new FST-based method to uncover local adaptation using environmental variables. Methods in Ecology and Evolution 6, 1248–1258 (2015).

21. Gautier, M. Genome-wide scan for adaptive divergence and association with population specific covariates. Genetics 201, 1555–1579 (2015).

22. Patterson, N. et al. Ancient admixture in human history. Genetics 192, 1065–1093 (2012).

23. Pickrell, J. K. & Pritchard, J. K. Inference of population splits and mixtures from genome-wide allele frequency data. PLos genetics 8, e1002967 (2012).

24. Racimo, F., Berg, J. J. & Pickrell, J. K. Detecting polygenic adaptation in admixture graphs. Genetics, genetics–300489 (2018).

25. Nicholson, G. et al. Assessing population differentiation and isolation from single-nucleotide polymorphism data. Journal Of The Royal Statistical Society: Series B (Statistical Methodology) 64, 695–715 (2002).

26. Skoglund, P. et al. Reconstructing prehistoric African population structure. Cell 171, 59–71 (2017).

27. Leppälä, K., Nielsen, S. V. & Mailund, T. admixturegraph: an R package for admixture graph manipulation and fitting. Bioinformatics 33, 1738–1740 (2017).

28. Lipson, M. et al. Efficient moment-based inference of admixture parameters and sources of gene flow. Molecular biology and Evolution 30, 1788–1802 (2013).

29. Haller, B. C. & Messer, P. W. SLiM 2: Flexible, interactive forward genetic simulations. Molecular biology and evolution 34, 230–240 (2016).

30. Consortium, I. G. P. et al. A global reference for human genetic variation. Nature 526, 68 (2015).

31. Lazaridis, I. et al. Ancient human genomes suggest three ancestral populations for present-day Europeans. Nature 513, 409 (2014).

32. Delaneau, O., Howie, B., Cox, A. J., Zagury, J.-F. & Marchini, J. Haplotype estimation using sequencing reads. The American Journal of Human Genetics 93, 687–696 (2013).

33. Das, S. et al. Next-generation genotype imputation service and methods. Nature genetics 48, 1284 (2016).

34. Durinck, S., Spellman, P. T., Birney, E. & Huber, W. Mapping identifiers for the integration of genomic datasets with the R/Bioconductor package biomaRt. Nature protocols 4, 1184 (2009).

35. Upadhyay, M. et al. Genetic origin, admixture and population history of aurochs (Bosprimigenius) and primitive European cattle. Heredity 118, 169 (2017).

36. Kim, J. et al. The genome landscape of indigenous African cattle. Genome biology 18, 34 (2017).

37. Da Fonseca, R. et al. Consequences of breed formation on patterns of genomic diversity and differentiation: the case of highly diverse peripheral Iberian cattle (submitted).

38. Elsik, C. G., Tellam, R. L., Worley, K. C., et al. The genome sequence of taurine cattle: a window to ruminant biology and evolution. Science 324, 522–528 (2009).

39. Nicolazzi, E. L. et al. SNPchiMp v. 3: integrating and standardizing single nucleotide polymorphism data for livestock species. BMC genomics 16, 283 (2015).

40. Korneliussen, T. S., Albrechtsen, A. & Nielsen, R. ANGSD: analysis of next generation sequencing data. BMC Bioinformatics 15, 356 (2014).

41. Árnason, E. & Halldórsdóttir, K. Nucleotide Variation and Balancing Selection at the *Ckma* gene in Atlantic cod: analysis with multiple merger coalescent models. PeerJ 3, e786 (2015).

42. ICES. *Spawning and life history information for North Atlantic cod stocks* ICES Cooperative Research Report 274 (International Council for the Exploration of the Sea, Copenhagen, Denmark, 2005). <http://www.ices.dk.

43. Jakobsson, J. & for the Exploration of the Sea, I. C. Cod and Climate Change: Proceedings of a Symposium Held in ReykjavíK, 23-27 August 1993 <https://books.google.is/books?id=lzkcAQAAIAAJ (International Council for the Exploration of the Sea, 1994).

44. Tørresen, O.K. et al. An Improved Genome Assembly Uncovers Prolific Tandem Repeats in Atlantic cod. BMC Genomics 18. doi:10.1186/s12864-016-3448-x. <https://doi.org/10.1186/s12864-016-3448-x> (Jan. 2017).

45. Li, H. & Durbin, R. Fast and Accurate Short Read Alignment with Burrows-Wheeler Transform. Bioinformatics 25, 1754–1760 (May 2009).

46. McKenna, A. et al. The Genome Analysis Toolkit: A MapReduce framework for analyzing next-generation DNA sequencing data. Genome Research 20, 1297–1303 (July 2010).

47. DePristo, M. A. et al. A Framework for Variation Discovery and Genotyping Using Next-Generation DNA sequencing data. Nature Genetics 43, 491–498 (Apr. 2011).

48. Halldórsdóttir, K. & Árnason, E. Whole-genome sequencing uncovers cryptic and hybrid species among Atlantic and Pacific cod-fish. BioRxiv, 034926 (2015).

49. Árnason, E. & Halldórsdóttir, K. Codweb: Whole-Genome Sequencing Uncovers Extensive Reticulations Fueling Adaptation Among Atlantic, Arctic, and Pacific Gadids. In Press (2018).

50. Li, H. Improving SNP discovery by base alignment quality. Bioinformatics 27, 1157–1158 (2011).

51. Sugden, L. A. et al. Localization of adaptive variants in human genomes using averaged one-dependence estimation. Nature communications 9, 703 (2018).

52. Kern, A. D. & Schrider, D. R. diploS/HIC: an updated approach to classifying selective sweeps. G3: Genes, Genomes, Genetics, g3–200262 (2018).

53. Akbari, A. et al. Identifying the favored mutation in a positive selective sweep. Nature methods (2018).

54. Mathieson, I. et al. Genome-wide patterns of selection in 230 ancient Eurasians. Nature 528, 499 (2015).

55. Bersaglieri, T. et al. Genetic signatures of strong recent positive selection at the lactase gene. The American Journal of Human Genetics 74, 1111–1120 (2004).

56. Grossman, S. R. et al. Identifying recent adaptations in large-scale genomic data. Cell 152, 703–713 (2013).

57. Racimo, F., Marnetto, D. & Huerta-Sanchez, E. Signatures of archaic adaptive introgression in present-day human populations. Molecular biology and evolution 34, 296–317 (2016).

58. Vernot, B. & Akey, J. M. Resurrecting surviving Neandertal lineages from modern human genomes. Science, 1245938 (2014).

59. Ohashi, J., Naka, I. & Tsuchiya, N. The impact of natural selection on an ABCC11 SNP determining earwax type. Molecular biology and evolution 28, 849–857 (2010).

60. Liu, S. et al. Low Pass Genomes of 141,431 Chinese Reveal Patterns of Viral Infection, Novel Phenotypic Associations, and the Genetic History of China (2018).

61. Andrés, A. M. et al. Targets of balancing selection in the human genome. Molecular biology and evolution 26, 2755–2764 (2009).

62. Stella, A., Ajmone-Marsan, P., Lazzari, B. & Boettcher, P. Identification of selection signatures in cattle breeds selected for dairy production. Genetics (2010).

63. De Simoni Gouveia, J. J. et al. Genome-wide search for signatures of selection in three major Brazilian locally adapted sheep breeds. Livestock Science 197, 36–45 (2017).

64. Mastrangelo, S. et al. Genome-wide scan of fat-tail sheep identifies signals of selection for fat deposition and adaptation. Animal Production Science (2018).

65. Keane, O. M. et al. Gene expression profiling of naive sheep genetically resistant and susceptible to gastrointestinal nematodes. BMC genomics 7, 42 (2006).

66. Ohtsuka, M. et al. NFAM1, an immunoreceptor tyrosine-based activation motif-bearing molecule that regulates B cell development and signaling. Proceedings of the National Academy of Sciences 101, 8126–8131 (2004).

67. Bahbahani, H., Afana, A. & Wragg, D. Genomic signatures of adaptive introgression and environmental adaptation in the Sheko cattle of southwest Ethiopia. Plos one 13, e0202479 (2018).

68. Plyte, S. E., Hughes, K., Nikolakaki, E., Pulverer, B. J. & Woodgett, J. R. Glycogen synthase kinase-3: functions in oncogenesis and development. Biochimica et Biophysica Acta (BBA) Reviews on Cancer 1114, 147–162 (1992).

69. Schlüter, G., Kremling, H. & Engel, W. The gene for human transition protein 2: nucleotide sequence, assignment to the protamine gene cluster, and evidence for its low expression. Genomics 14, 377–383 (1992).

70. Engel, W., Keime, S., Kremling, H., Hameister, H. & Schlüter, G. The genes for protamine 1 and 2 (PRM1 and PRM2) and transition protein 2 (TNP2) are closely linked in the mammalian genome. Cytogenetic and Genome Research 61, 158–159 (1992).

71. Litwack, E. D., Babey, R., Buser, R., Gesemann, M. & O’leary, D. D. Identification and characterization of two novel brain-derived immunoglobulin superfamily members with a unique structural organization. Molecular and Cellular Neuroscience 25, 263–274 (2004).

72. Randhawa, I. A., Khatkar, M. S., Thomson, P. C. & Raadsma, H. W. A meta-assembly of selection signatures in cattle. PLos one 11, e0153013 (2016).

73. Shaheen, R. et al. A founder CEP120 mutation in Jeune asphyxiating thoracic dystrophy expands the role of centriolar proteins in skeletal ciliopathies. Human molecular genetics 24, 1410–1419 (2014).

74. Davis, C. A. et al. PRISM/PRDM6, a transcriptional repressor that promotes the proliferative gene program in smooth muscle cells. Molecular and cellular biology 26, 2626–2636 (2006).

75. Chen, W. V. & Maniatis, T. Clustered protocadherins. Development 140, 3297–3302 (2013).

76. Hayashi, S. & Takeichi, M. Emerging roles of protocadherins: from self-avoidance to enhancement of motility. J Cell Sci, jcs–166306 (2015).

77. Fukuda, E. et al. Down-regulation of protocadherin-*α* A isoforms in mice changes contextual fear conditioning and spatial working memory. European Journal of Neuroscience 28, 1362–1376 (2008).

78. Montague, M. J. et al. Comparative analysis of the domestic cat genome reveals genetic signatures underlying feline biology and domestication. Proceedings of the National Academy Of Sciences 111, 17230–17235 (2014).

79. Wang, X. et al. Genomic responses to selection for tame/aggressive behaviors in the silver fox (Vulpes vulpes). Proceedings of the National Academy of Sciences 115, 10398–10403 (2018).

80. Lee, H.-J. et al. Deciphering the genetic blueprint behind Holstein milk proteins and production. Genome biology and evolution 6, 1366–1374 (2014).

81. Peng, C. et al. Ablation of vacuole protein sorting 18 (Vps18) gene leads to neurodegeneration and impaired neuronal migration by disrupting multiple vesicle transport pathways to lysosomes. Journal of Biological Chemistry 287, 32861–32873 (2012).

82. Yang, J. et al. Differential expression of genes in milk of dairy cattle during lactation. Animal genetics 47, 174–180 (2016).

83. Hemmer-Hansen, J. et al. A genomic island linked to ecotype divergence in A tlantic cod. Molecular Ecology 22, 2653–2667 (2013).

84. Bradbury, I. R. et al. Genomic islands of divergence and their consequences for the resolution of spatial structure in an exploited marine fish. Evolutionary Applications 6, 450–461 (2013).

85. Kirubakaran, T. G. et al. Two adjacent inversions maintain genomic differentiation between migratory and stationary ecotypes of Atlantic cod. Molecular ecology 25, 2130–2143 (2016).

86. Lee, K. M. & Coop, G. Distinguishing among modes of convergent adaptation using population genomic data. Genetics, genetics–300417 (2017).

87. Bongiorni, S. et al. Skeletal muscle transcriptional profiles in two Italian beef breeds, Chianina and Maremmana, reveal breed specific variation. Molecular biology reports 43, 253–268 (2016).

88. Thorsteinsson, V., Pálsson, Ó. K., Tómasson, G. G., Jónsdóttir, I. G. & Pampoulie, C. Consistency in the behaviour types of the Atlantic cod: repeatability, timing of migration and geo-location. Marine Ecology Progress Series 462, 251–260 (2012).

89. Pálsson, Ó. K. & Thorsteinsson, V. Migration patterns, ambient temperature, and growth of Icelandic cod (Gadus morhua): evidence from storage tag data. Canadian Journal of Fisheries And Aquatic Sciences 60, 1409–1423 (2003).

90. Pampoulie, C., Jakobsdóttir, K. B., Marteinsdóttir, G. & Thorsteinsson, V. Are vertical behaviour patterns related to the pantophysin locus in the Atlantic cod (Gadus morhua L.)? Behavior genetics 38, 76–81 (2008).

91. Pampoulie, C. et al. Rhodopsin gene polymorphism associated with divergent light environments in Atlantic cod. Behavior genetics 45, 236–244 (2015).

92. Pogson, G. H. Nucleotide polymorphism and natural selection at the pantophysin (Pan I) locus in the Atlantic cod, Gadus morhua (L.) Genetics 157, 317–330 (2001).

93. Sodeland, M. et al. Islands of Divergence in the Atlantic cod genome represent polymorphic chromosomal rearrangements. Genome biology and evolution 8, 1012–1022 (2016).

94. Berg, P. R. et al. Trans-oceanic genomic divergence of Atlantic cod ecotypes is associated with large inversions. Heredity 119, 418 (2017).

95. Barth, J. M. et al. Genome architecture enables local adaptation of Atlantic cod despite high connectivity. Molecular ecology 26, 4452–4466 (2017).

96. Hahne, F. & Ivanek, R. in Statistical Genomics 335–351 (Springer, 2016).

